# Single-cell atlas of mouse limb development reveals a complex spatiotemporal dynamics of skeleton formation

**DOI:** 10.1101/2022.05.07.490557

**Authors:** Svetlana Markman, Mor Zada, Eyal David, Amir Giladi, Ido Amit, Elazar Zelzer

**Affiliations:** Department of Molecular Genetics, Weizmann Institute of Science, Rehovot, Israel; Department of Immunology, Weizmann Institute of Science, Rehovot, Israel

## Abstract

Limb development has long served as a model system for coordinated spatial patterning of progenitor cells. Here, we identify a population of naïve limb progenitors and show that they differentiate progressively to form the skeleton in a complex nonconsecutive three-dimensional pattern.

Single-cell RNA sequencing of the developing mouse forelimb revealed three progenitor states: naïve, proximal and autopodial, as well as *Msx1* as a marker for the naïve progenitors. In vivo lineage tracing confirmed this role and localized the naïve progenitors to the outer margin of the limb, along the anterior-posterior axis. Sequential pulse-chase experiments showed that the progressive transition of *Msx1^+^* naïve progenitors into proximal and autopodial progenitors coincides with their differentiation to *Sox9^+^* chondroprogenitors, which occurs along all the forming skeletal segments.

Indeed, tracking the spatiotemporal sequence of differentiation showed that the skeleton forms progressively in a complex pattern. These findings suggest a new model for limb skeleton development.

## Introduction

Limb development has long served as a central model system for studying organ formation. Over the years, extensive research has identified key components in the genetic program that controls both patterning and differentiation of the different tissues composing the limb, as well as the complex signaling involved in regulating this genetic program (Delgado and Torres, 2017; Huang, 2017; Johnson and Tabin, 1997; Nassari et al., 2017a; Niswander, 2003; Tickle, 2005; Zuniga, 2015). These studies have produced the basic concepts of how progenitor cells pattern and differentiate along the three axes to form a complex functional organ (Petit et al., 2017; Tabin and Wolpert, 2007a; Zeller et al., 2009).

The mouse forelimb starts to develop around E9.5 as a small outgrowth from the body wall. Initially, the limb bud comprises seemingly homogeneous undifferentiated mesenchymal cells covered by a layer of ectoderm. At the distal side, along the anterior-posterior border, the ectoderm thickens to form the apical ectodermal ridge (AER). As development proceeds, the limb is elongated and the hand plate is formed. Concurrently, the development of the limb skeleton is initiated, as mesenchymal cells form condensations that prefigure the future skeletal elements.

The vertebrate limb skeleton is organized in three segments: stylopod, containing humerus in the forelimb or femur in the hindlimb, zeugopod, comprising radius and ulna or tibia and fibula and autopod, comprising the wrist or ankle and digits. Surgical removal of the AER during early wing-bud development resulted in severe truncation of the distal elements. Moreover, the later the AER was removed, the more distal elements were formed. These findings led to the perception that the skeletal elements of the limb form in a proximal-to-distal order under the regulation of the AER (Saunders, 1948; Summerbell et al., 1973). Several models have attempted to explain this mode of development (Tabin and Wolpert, 2007a). The progress zone model is named after a distal domain under the AER, where limb progenitor cells are postulated to be located (Saunders, 1948; Summerbell et al., 1973). According to this model, the longer these progenitors spend in the progress zone, the more distal their progeny become. Once the cells exit this domain, their fate is determined. The result is that the first cells to leave the progress zone form the stylopod, the next to leave form the zeugopod, and the cells that exit last form the autopod (Saunders, 1948; Summerbell et al., 1973; Wolpert, 2002). An alternative model posits that progenitors of the limb segments are specified early in development, organized in three parallel stripes, and then expand progressively in a proximodistal order (Dudley et al., 2002; Sun et al., 2002). The two-signal model suggests that proximal and autopod progenitors are specified by two opposing signals deriving from the flank and AER, respectively. Later on, as the limb bud grows, a third domain of the zeugopod is formed in the middle (Mariani et al., 2008; Mercader et al., 1999a, 2000).

Recently, several works have studied mouse and chick limb development at single-cell resolution (Desanlis et al., 2020a; Feregrino et al., 2019; Kelly et al., 2020). Despite their findings, fundamental aspects of this process are still missing. For example, the identity of limb progenitors and their spatial distribution are unclear, as we lack marker genes to identify them. The temporal changes the transcriptome of these progenitors undergo during development and the sequence by which they differentiate to form the limb skeleton have yet to be uncovered, too.

In this work, we establish a single-cell atlas of the developing limb and characterize limb progenitors. We identified three progenitor populations, namely naïve, proximal and autopodial limb progenitors. We established *Msx1* as a marker for the naïve progenitors and their location in the outer margin of the developing limb, along the anterior-posterior axis. We then showed that these *Msx1^+^* naïve progenitors transit progressively and simultaneously into either proximal or autopodial progenitors. Moreover, the progressive contribution of these progenitors to the forming skeleton occurs simultaneously all along the proximal-distal axis of the limb. Finally, temporal analysis of the differentiation of *Msx1* lineage cells revealed that the skeleton forms in a complex nonconsecutive three-dimensional pattern, which extends to the level of the single element.

## Results

### Single-cell RNA sequencing provides a comprehensive cellular and molecular atlas of the major mesenchyme-derived cells types of the developing limb

To date, the identity of limb mesenchymal progenitor cells and their differentiation paths are only partially understood. To obtain a deeper and unbiased molecular characterization of mesenchymal progenitor cells, we generated transcriptional maps of mesenchymal lineages in the developing limb between E10.5 and E14.5 by applying a massively parallel single-cell RNA-seq (MARS-seq). During this time window, mesenchymal cells undergo major patterning and differentiation to form the different tissues of the limb, including muscle connective tissue, tendons, ligaments and skeleton. To ensure representation of the different cell types and differentiation states, including rare subpopulations, we combined lineage and reporter-based single-cell analysis using *Sox9* and *Scx*, the earliest known markers for skeletal and tendon cells, respectively (Akiyama et al., 2002a; Bi et al., 1999; Schweitzer et al., 2001).

At E10.5, we sampled a *Sox9-GFP* transgenic mouse line and collected both *Sox9^+^* and negative cell populations. Because *Sox9* marks multiple cell types (Akiyama‡ et al., 2005; Nagakura et al.; Soeda et al., 2010; Sugimoto et al., 2013), to follow the dynamics of different lineages and to ensure the representation of tendons we generated a compound mouse model containing *Sox9-CreER^T2^* (Soeda et al., 2010)*, tdTomato* (Madisen et al., 2010) and *Scx-GFP* (Pryce et al., 2007). Tamoxifen was administered at different developmental time points between E9.5 and E12.5 and samples were harvested 48 h later. Thus, we sampled four cell populations: tdTomato-positive (*Sox9^+^* skeletal lineage), GFP*-* positive (*Scx^+^* tendon cells), tdTomato-GFP double-positive cells, and double-negative cells (other progenitors). Then, 32,000 quality-filtered cells (see Methods) were subjected to MARS-seq to generate a limb cellular atlas. We used the MetaCell algorithm (Baran et al., 2019) to identify homogeneous and robust groups of cells, referred to as meta-cells (MCs; see Methods). To focus on mesenchymal cell lineages, MCs of non-mesenchymal origin, such as red blood cells, muscle cells, ectoderm-derived cells and Schwann cells, were excluded from the analysis (Table S1; Methods). The remaining 250 MCs were grouped into 12 molecularly distinct populations (Fig. 1B-F; Fig. S1).

**Figure 1.**
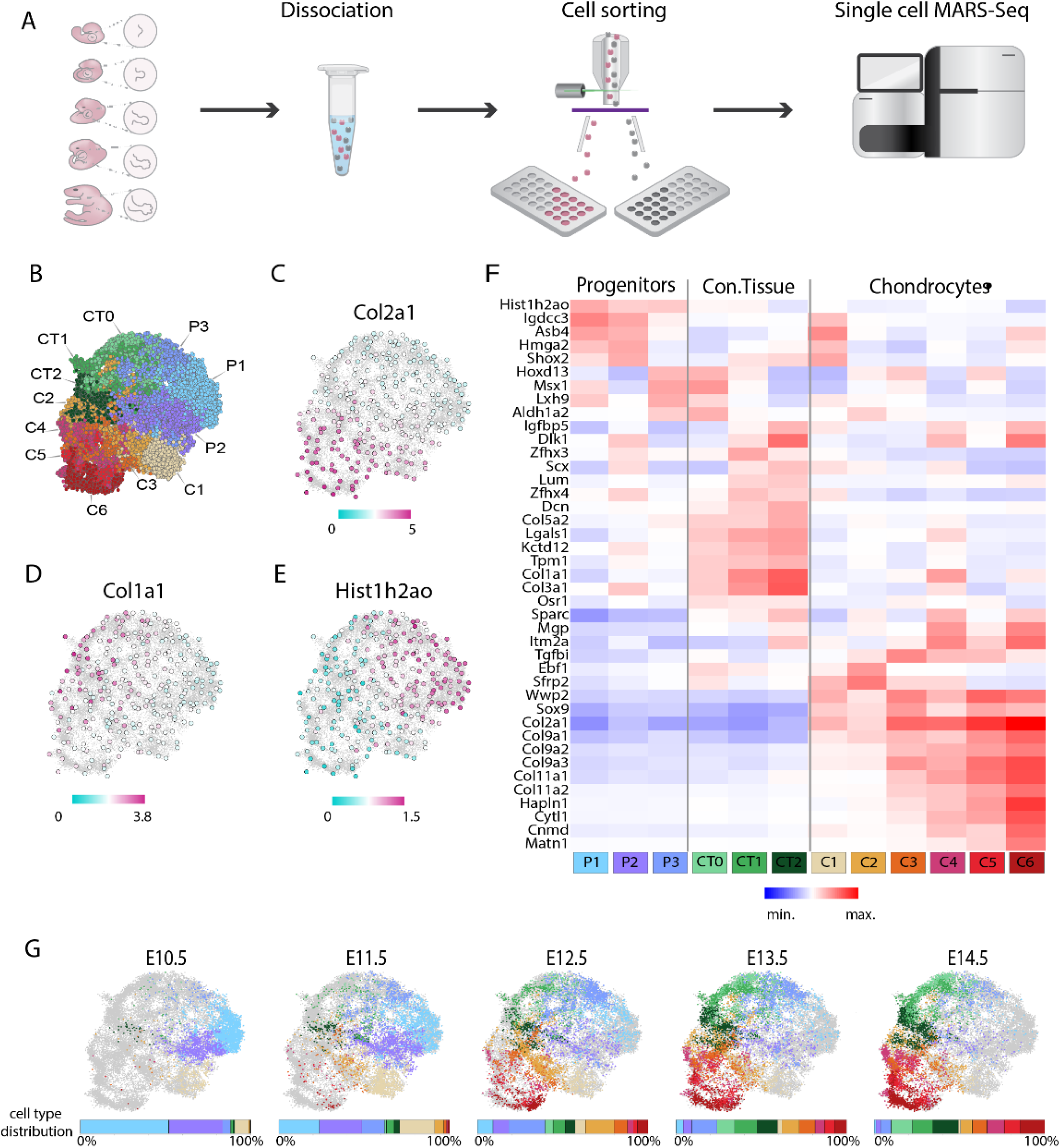
Single-cell RNA sequencing of mouse forelimb mesenchymal cells during embryonic development. (A) Scheme showing the experimental design. Forelimbs were disassociated into single cells, which were then FACS-sorted into 384-well plates. Isolated cells were subjected to massively parallel single-cell RNA sequencing (MARS-seq). At E10.5, *Sox9-GFP* embryos (n=3) were used to separate between GFP-positive and negative cells. Additionally, cells were collected from *Sox9-CreER^T2^; tdTomato;Scx-GFP* embryos (n=2) without Cre activation. For collection of cells from E11.5-E14.5 embryo forelimbs, we used *Sox9-CreER^T2^;tdTomato;ScxGFP* mice (n=4-5 per stage), in which *Sox9*^+^ cells were labeled by tamoxifen administration 48 h before harvesting and sorted into tdTomato^+^GFP^-^, tdTomato^-^GFP^+^, double-positive and double-negative cells. (B) *k*-nearest neighbors (*k*-NN) graph of 26,131 mesenchymal cells (250 metacells) associated with 12 annotated and color-coded cell types and states. (C-E) The 250 metacells were subdivided into three main cell populations, as shown by log_2_ fold change in gene expression of *Col2a1* (C) *Col1a1* (D) and *Hist1h2ao* (C) genes projected onto the *k*-NN graph. (F) Heatmap showing log_2_ fold change in expression of known and newly identified marker genes in 12 transcriptionally distinct cell populations. (G) Projection of single cells onto the *k*-NN graph shows cell type distribution at different developmental stages.

To annotate clusters as chondrocytes or connective tissue fibroblasts, we performed differential gene expression analysis. We used the established markers *Sox9*, *Col2a*, *Wwp2*, and *Col9a2* for the former and *Col1a1*, *Col3a1*, *Scx*, *Osr1*, *Dcn*, and *Lum* for the latter (Akiyama et al., 2002b; Bell et al., 1997; Bi et al., 1999; Lefebvre et al., 1997; Liu et al., 2015; Nakamura et al., 2011; Schweitzer et al., 2001; Stricker et al., 2012; Vallecillo-García et al., 2017). Results showed that subsets C1-6 represent distinct chondrocyte subpopulations and groups CT0-CT2 represent distinct connective tissue fibroblast subpopulations (Fig. 1C,D,F; Fig. S1). Cells in subsets P1 and P3 expressed neither fibroblast nor chondrocyte markers and were therefore annotated as mesenchymal progenitors (Fig. 1C-F). A large gene signature defined these progenitor types including *Hist1h2ao* (Table S2), which was expressed by P1-P3 cells but not by any of the differentiated cell types (Fig. 1E,F). P1 cluster was characterized by co-expression of *Hmga2*, *Asb4*, *Igdcc3*, *Msx1* and *Lhx9*. *Msx1* and *Lhx9* are transcription factors (TFs) that are expressed in limb mesenchyme and play a key role in limb patterning (Bensoussan-Trigano et al., 2011; Lallemand et al., 2005; Tzchori et al., 2009), whereas *Hmga2* is widely expressed in undifferentiated cells during embryogenesis (Ulrike Hirning-Folz, Monika Wilda, Volkhard Rippe, Jorn Bullerdiek, 1998; Xianjin Zhou, Kathleen F. Benson, 1995). *Asb4* is a ubiquitin ligase and *Igdcc3* is a member of the immunoglobulin superfamily, with a suggested role in early embryogenesis (Salbaum, 1998). Interestingly, these progenitors lacked a spatial signature.

P2 cells expressed a gene module that largely overlapped with that of P1; however, P2 cells lacked the expression of *Msx1* and *Lxh9*. Instead, this cluster was characterized by high expression of *Shox2*, *Dlk1*, *Zfhx3-4*, *Scx*, and *Col3a1*. *Shox2* is a TF that is expressed at the proximal limb bud, where it acts as a patterning gene (Sun et al., 2013). The P3 subset was characterized by high expression of *Hoxd13*, *Msx1*, *Lhx9* and *Aldh1a2*. The known role of these genes in regulating autopod patterning suggests that this cluster represents progenitors of the autopod (Fromental-Ramain et al., 1996; Scotti et al., 2015a). Overall, we identified three progenitor populations, two of them carrying a signature of either proximal or autopodial markers, whereas the third population lacked a typical signature. Subsets CT0-CT2 were characterized by the expression of *Col5a1*, *Col3a1*, *Col1a1* and *Osr1*, all of which are markers of connective tissue/tendon. In addition, these cells also expressed *Lgals1*, which was implicated in modulating cell-cell and cell-matrix interactions. Other identified markers were *Kctd12*, encoding for a potassium channel, and tropomyosin (*Tpm1*). Interestingly, CT0 cells did not express the key tendon marker *Scx*. Instead, this cluster was characterized by expression of the markers *Msx1*, *Hoxd13*, and *Aldh1a2*, similar to the P3 signature. The transcriptional similarity between CT0 and P3 suggests that some P3 autopod progenitors give rise to CT0 cells. CT1 and CT2 were both characterized by the expression of tendon markers *Scx*, *Dcn* and *Lum* (Liu et al., 2015; Schweitzer et al., 2001). In addition, cells of both clusters expressed the TF *Zfhx4*, *Zfhx3* marked CT1 cells, whereas CT2 was marked by high expression of *Dlk1*, *Igfbp5* and *Sparc.* In the chondrocyte compartment, subsets C1 and C2 displayed low expression levels of cartilage-specific ECM genes (cECM), such as *Col2a1*, *Col9a1-3*, *Col11a1-2* and *Hapln1*, indicating the early differentiation stage of these clusters. In line with this, C1 displayed high expression levels of *Asb4*, *Shox2*, *Hmga2* and *Igdcc3*, which marked cluster P2, suggesting that cluster C1 originates from P2 cells. Cluster C2 displayed high expression of the TFs *Ebf1* and *Sfrp2*, a soluble modulator of Wnt signaling, and of the autopod marker *Hoxd13*, suggesting that this cluster represents early autopodial chondrocytes. Subset C3 displayed expression of *Hoxd13*, intermediate levels of cECM genes, and high *Tgfbi* expression, suggesting that it represents more mature autopodial chondrocytes. Subset C4 displayed intermediate expression levels of cECM genes in combination with several fibroblast markers, specifically high expression of *Col1a1* and low expression of *Scx*, *Col3a1* and *Lgals1*. These results suggest a chondro-tendinous identity of these cells (Blitz et al., 2013; Sugimoto et al., 2013). Subset C5 displayed high expression levels of cECM genes along with the autopod marker *Hoxd13*. Therefore, it represents the most mature state of autopodial chondrocytes. Subset C6 displayed the highest expression levels of all cECM genes, thus likely to represent the most differentiated chondrocytes.

To elucidate the temporal dynamics of identified cell populations, we annotated the enrichment of each population over time. As shown in Figure 1G, dramatic changes in cell type composition were observed during development. While P1 was the most abundant cell population at E10.5 (52%), it gradually decreased until it was completely diminished by E14.5. Proximal progenitors (P2) also constantly decreased from 31% at E10.5 to 3% by E14.5. In contrast, autopodial progenitors (P3) were rare at E10.5 (4%), increased up to 21% by E12.5, and later on started to decline reaching 6% by E14.5. Among the differentiated cell types, early proximal chondrocytes (C1) were rare at E10.5 (8%). By E11.5, C1 increased to 20% and early autopodial chondrocytes (C2) appeared (5%), concurrently with the appearance of tendon fibroblasts CT1 (4%) and CT2 (3%). At E12.5, C1 dramatically decreased (6%), while C2, CT1 and CT2 increased (16%, 7%, and 6%, respectively). Additionally, at E12.5 we saw the appearance of more mature chondrocytes C3-C6 (8%, 3%, 2%, and 6%, respectively) and CT0 (5%). At E13.5, C1 continued to decrease (2%) along with a decrease in C2 (10%), whereas more mature chondrocytes (C3-C6), CT1, CT2 and CT0 increased (chondrocytes, 10%, 5%, 6%, and 7%, respectively; CT, 13%, 12%, and 10%, respectively). By E14.5, C1 was diminished, C2 continued to decrease (7%), C3 and CT1 displayed a slight reduction (8%, 12%), while C4-C6, CT2 and CT0 further increased (to 10%, 9%, 14%, 13%, and 15%, respectively).

Overall, these data reveal the main mesenchyme-derived cells types in the developing limb, as well as valuable cell type-specific markers for studying their differentiation trajectories and dynamics. We identified three populations of progenitors, including autopodial and proximal progenitors and a third progenitor population that lacks spatial signature. Interestingly, we failed to identify a zeugopodial progenitor population. Finally, the gradual reduction in progenitor cells and increase in differentiated cells suggests a progressive differentiation process in the developing limb.

### Characterization of limb progenitors

Our meta-cell analysis identified three transcriptionally distinct populations of progenitors, namely P1, proximal P2 and autopodial P3 (Fig. 2A). To gain insight into transcriptional mechanisms and molecular pathways regulating these cells, we computationally extracted annotated MCs from these three subpopulations and computed the Pearson’s correlation coefficients for each pair of genes across all cells (see Methods). Hierarchical clustering of the correlation matrix (Fig. 2B; Table S3) revealed four gene modules. Module 1 was enriched for components of signaling pathways such as TGF-β/activin and BMP (*Bmp2*, *Gdf5*, *Dlx5*, *Dlx6*, *Inhba*, *Bambi*), Wnt (*Wnt5a*) and Fgf (*Fgf12*, *Sp9*), as well as for retinoic acid synthesis enzymes (*Aldh1a2*, *Rdh10*). This module also contained several TFs that regulate limb patterning (*Msx1*, *Msx2*, *Lhx2* and *Lxh9*) as well as TFs that are essential specifically for autopod patterning, such as *Hoxa13*, *Hoxd13*, *Hoxd12*, *Tfap2a*, and *Tfap2b* (Shen et al., 1997; Zhao et al., 2011), thus representing an autopodial genetic program. Module 2 was enriched with genes involved in matrix formation (*ccdc80*, *Lox*, *Eln*) and calcium binding proteins (*Egfl6*, *Sned1*, *Sparc*, *Piezo2*) as well as with Wnt signaling components *Dkk2* and *Fzd8*. These results suggest that this module represents a progressive stage of cell differentiation.

**Figure 2.**
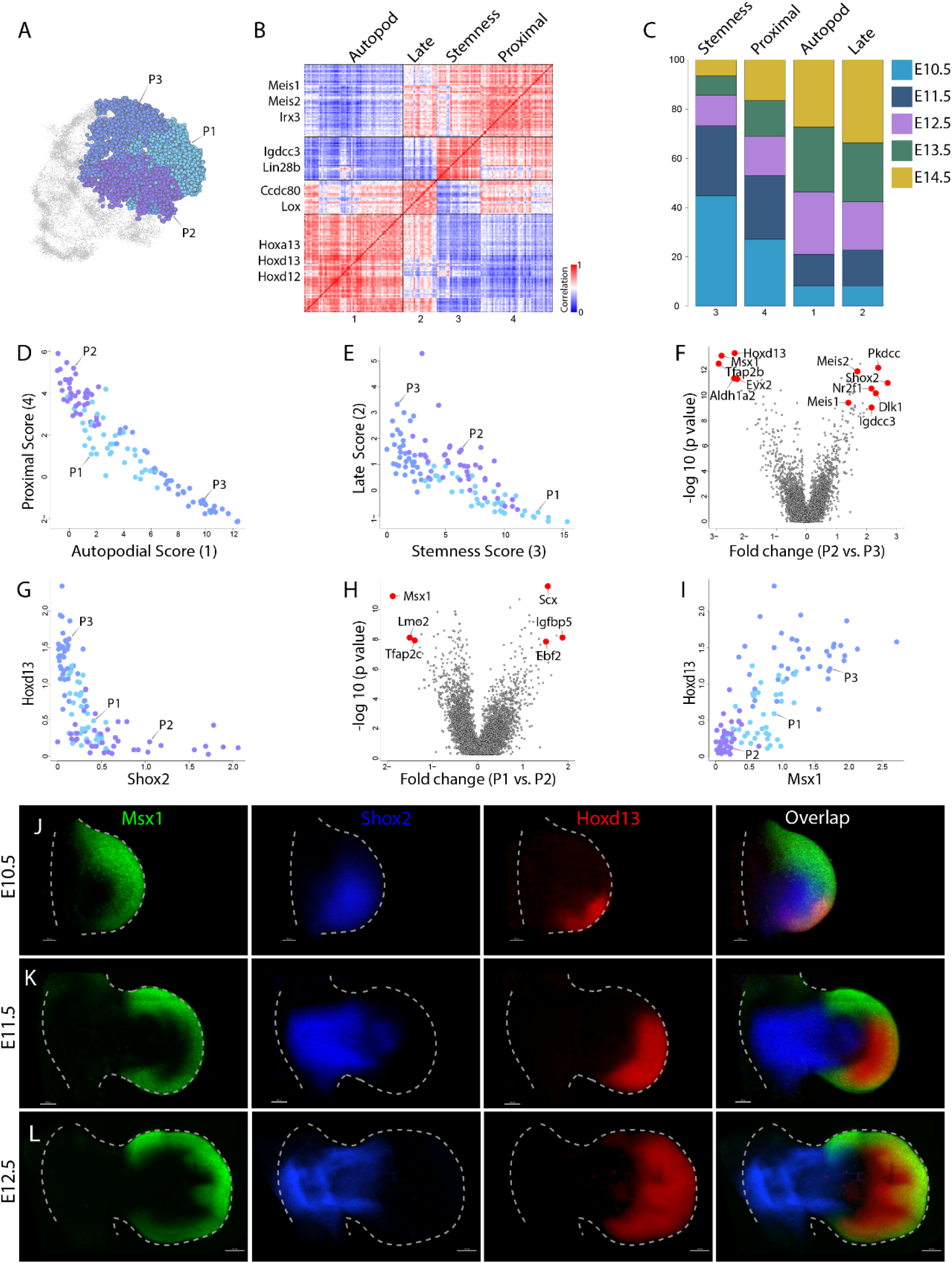
Characterization of mesenchymal progenitor cells. (A) *k*-NN graph of 10,241 progenitor cells, grouped into three subsets. Dots represent single cells, which were annotated and color-coded as in Figure 1B. (B) Heatmap showing hierarchical clustering of 175 genes that were most variably expressed by progenitor cells into four modules, based on gene-gene Pearson’s correlation. Representative genes are indicated for each gene module. (C) Graph showing the relative contributions of cells at various developmental stages to the total gene expression in each module. (D) Scatter plot showing the distribution of autopodial (*X*-axis) and proximal (*Y*-axis) scores in progenitor metacells. (E) Scatter plot showing the distribution of stemness (*X*-axis) and late (*Y*-axis) scores in progenitor meta-cells. (F) Volcano plot showing differentially expressed genes between P2 and P3 cells, presented as expression fold change (*X*-axis) and *p*-value (*Y*-axis, −log10 scale). The five most significantly differentially expressed genes and the two established proximal markers *Meis1* and *Meis2* are indicated by red dots. (G) Scatter plot showing the differences in *Shox2* (*X*-axis) and *Hoxd13* (*Y*-axis) expression across progenitor metacells.(H) Volcano plot showing differentially expressed genes between P1 and P2 cells, presented as expression fold change (*X*-axis) and *p*-value (*Y*-axis, −log_10_ scale). The three most significantly differentially expressed genes are indicated by red dots.(I) Scatter plot showing the differences in *Msx1* (*X*-axis) and *Hoxd13* (*Y*-axis) expression across progenitor metacells. (J-L) Maximum intensity projection (MIP) images of E10.5 (K), E11.5 (L) and E12.5 (M) whole-mount forelimbs stained for *Msx1* (green), *Shox2* (blue) and *Hoxd13* (red) mRNA using *in situ* HCR and imaged by light sheet microscopy. *Msx1* is expressed at the anterior-posterior margin of the limb in an arc-like pattern that is wider at the anterior side. *Hoxd13* expression expanded from a small dorsal posterior region at E10.5 to occupy most of the autopod at E12.5. *Shox2* is expressed in the proximal limb, stylopod and zeugopod. At each stage, n=2.

Module 3 was enriched for several signaling pathways, such as Wnt (*Rspo4*) and BMP (*Grem1*), and a subset of homeobox genes including *Hoxd4*, *Hoxd8*, *Hoxd9*, *Hoxa11*, and *Shox2*. Interestingly, this module was also enriched with genes associated with the maintenance of pluripotency and stem cell function, including *Igdcc3*, *Sall4*, *Lin28b*, *Tfapc2* and *Trim71*, suggesting that this module represents a stemness genetic program (Chang et al., 2012; Melton et al., 2010; Pastor et al., 2018; Patterson et al., 2011; Rybak et al., 2009; Wang et al., 2006; Worringer et al., 2014; Yu et al., 2007; Zhang et al., 2006, 2016; Zhao et al., 2011). Module 4 was enriched for signaling pathways such as Igf (*Igf2*, *Igfbp5*, *Igfbp3*, *Igf1*) and Wnt (*Ror1*, *Dact1*). Additionally, it contained proximally expressed genes such as *Meis1*, *Meis2*, *Pkdcc*, *Meox1*, *Pitx2*, *Emx2*, and *Irx3*, thus representing a proximal gene program (Campbell et al., 2012; Delgado et al., 2020; Li et al., 2014; Mercader et al., 1999b, 2009; Pellegrini et al., 2001; Probst et al., 2011; Reijntjes et al., 2007; Vickerman et al., 2011). Interestingly, module 4 also contained genes associated with tendon and connective tissue formation, such as *Scx*, *Tcf15*, *Osr2* and *Cxcl12* (Nassari et al., 2017b; Wilson-Rawls et al., 2004).

We next examined the temporal activity of these four modules (Fig. 2C). Results showed that 8% of the autopodial module 1 gene expression came from E10.5 cells, 12% from E11.5 cells and 25-27% from E12.5-E14.5 cells. These results suggest that autopodial gene program is detectable already at E10.5 cells and becomes more prominent at E12.5-E14.5. In module 2, 8% of the gene expression was associated to E10.5 cells, with constantly increasing contribution at later developmental stages (15% at E11.5, 19% at E12.5, 24% at E13.5, and 34% at E14.5), suggesting that this module represents a late genetic program. In the stemness module 3, 45% of gene expression was associated to E10.5 cells, followed by a decline in contribution at later stages (28% at E11.5, 12% at E12.5, 8% at E13.5, and 6% at E14.5). These results indicate that cells turn off the expression of stemness genes as development proceeds. In the proximal module 4, gene expression was associated mostly to E10.5 and E11.5 cells (27% and 26%, respectively), whereas E12.5-E14.5 cells contributed 16%, 14%, and 16%, respectively. These results suggest that proximal genes are expressed by cells throughout this period.

Next, we utilized our gene module information to classify the three identified progenitor populations. To determine how these functional gene modules are distributed across cell types, we calculated scores of each module for each MC. Examination of proximal (4) vs autopodial (1) module scores showed that MCs of proximal P2 and autopodial P3 were completely separated (Fig. 2D). P2 was enriched with proximal module genes, whereas P3 was enriched with autopodial module genes, further confirming our annotation. Interestingly, P1 MCs had low levels of both proximal and autopodial scores, with some MCs overlapping with proximal or autopodial MCs. The location of P1 MCs between P2 and P3 with some overlap suggests that P1 MCs transit into proximal and autopodial progenitor states.

P1 MCs had higher levels of expression of the stemness module (3) scores, as compared to P2 and P3, which expressed intermediate and high expression of the late module (1) scores, respectively (Fig. 2E), further supporting the naïve state of P1. Finally, examination of proximal and stemness scores showed a clear separation between the three progenitor groups. P1 was characterized by high-to-intermediate stemness score combined with low proximal scores, whereas P2 was characterized by intermediate stemness scores combined with high proximal scores. P3 displayed the lowest levels of both proximal and stemness scores (Fig. S2A,B).

Overall, these results classify limb progenitors as proximal, autopodial and naïve, and uncover TFs and signaling circuits that regulate these populations. Temporally, we show that proximal and autopodial gene modules are active throughout the developmental process. Additionally, a transition from stemness to late genetic program was observed. To reveal the spatial distribution of these progenitors *in vivo*, we searched for specific markers for the three progenitor populations. For that, we first analyzed the most differentially expressed genes between proximal and autopodial progenitors (Table S4). As shown in Fig. 2F,G and Table S4, *Shox2* and the known proximal markers *Meis1* and *Meis2* were upregulated in proximal progenitors, while *Tfap2b, Hoxd13, Hoxa13* and *Msx1* were among the most differentially expressed genes in the autopodial progenitors. We selected *Shox2* and *Hoxd13* as markers for our *in situ* experiments due to the higher expression levels of these two genes. Naïve progenitors displayed significantly lower levels of *Hoxd13* (1.3 fold, p = 10.7 –log_10_; Table S5). Comparison of gene expression between naïve and proximal subpopulations showed that *Msx1* was among the most differentially expressed genes by P1 cells (Fig. 2H; Table S6). Because *Msx1* was also expressed by P3 cells, we examined the combination of *Msx1* and *Hoxd13*. As seen in Fig. 2I, this combination clearly separated between P1 and P3 MCs. Thus, we defined *Msx1^+^/Hoxd13^-^* cells as naïve progenitors, *Shox2^+^* cells as proximal progenitors and *Msx1^+^/Hoxd13^+^* as autopodial progenitors.

To study the spatiotemporal distribution of the three progenitor populations during limb development, we conducted whole-limb triple *in situ* hybridization chain reaction (HCR) using *Msx1*, *Shox2* and *Hoxd13* probes. As seen in Fig. 2J, at E10.5, *Msx1* was expressed in the outer margin of the limb forming an arc-like pattern along the anterior-posterior axis. The arc extended both dorsally and ventrally from the AP midline, more so dorsally (Fig. S2C-E). At the anterior-proximal side of the arc, *Msx1* expression domain was the widest (Fig. 2J, Fig. S2F,G). At the dorsal side, the most posterior *Msx1* expression domain overlapped with *Hoxd13* expression domain, demarcating the location of the autopodial progenitors (Fig. 2J, Fig. S2H,I). *Shox2* proximal progenitors were found at the core of the limb bud, encircled by the *Msx1* expression domain.

At E11.5 (Fig. 2K, Fig. S2J-P), the arc-like pattern of *Msx1* expression along the AP axis was maintained, as were the size asymmetries along the AP and DV axes. The overlap between *Msx1* and *Hoxd13* expression domains expanded dorsally and ventrally as well as anteriorly, occupying the most distal front of the limb. *Shox2* expression domain extended throughout most of the proximal limb segment.

At E12.5 (Fig. 2L), *Msx1* arc-like expression domain was still visible. *Msx1* and *Hoxd13* expression domains occupied the interdigital space and most of the outer margin of the autopod, with the exception of the anterior region of the developing thumb and a small posterior region, which were positive only for *Msx1* (Fig. S2Q-R). *Shox2* expression domain occupied most of the proximal limb segment. Areas of overlap between *Shox2* and *Msx1* and between *Msx1* and *Hoxd13* were observed for several days, supporting the transition of P1 cells into either P2 or P3 cells, as suggested by our analysis.

Together, these data suggest that P1 represents naïve multipotent progenitor cells, which are marked by *Msx1* and are located in the outer margin of the forming limb. These cells differentiate into P2 proximal progenitors, marked by *Shox2*, and P3 autopodial progenitors, which are co-marked by *Msx1* and *Hoxd13*. Another important finding is the co-existence of P2 and P3 cells at E10.5-E12.5, which suggests that during this time window, P1 cells transit progressively and simultaneously into P2 and P3 cells.

### *Msx1* marks the naïve progenitors of the limb

A central hypothesis raised by our single-cell data is that the transcription factor *Msx1* marks the most naïve limb mesenchymal progenitors. If indeed this is the case, this TF should be expressed at the onset of limb development and its lineage should give rise to all mesenchyme-derived tissues, including cartilage, tendon and muscle connective tissue. To test this prediction, we first studied the expression of *Msx1* at E9.5, the onset of limb development. As seen in Figure 3A and in agreement with previous studies (Coudert et al., 2005), *Msx1* expression was observed in the cells of the forming forelimb. To examine directly the contribution of the *Msx1* lineage to the different mesenchymal limb tissues, we utilized the previously described *Msx1-CreER^T2^* knock-in mouse line (Lallemand et al., 2013) crossed with *Rosa26-tdTomato* (Madisen et al., 2010) and *Scx-GFP* (Pryce et al., 2007) mice. As seen in Figure 3B-F, a single dose of tamoxifen at E9.5 marked cells of the entire skeleton, tendons and muscle connective tissue in the E14.5 forelimb. These results confirm our single-cell data showing that *Msx1* marks the naïve multipotent mesenchymal progenitors. Moreover, they confirm the high efficiency of the *Msx1-CreER^T2^* knock-in allele in activating the *Rosa26-tdTomato* reporter.

**Figure 3.**
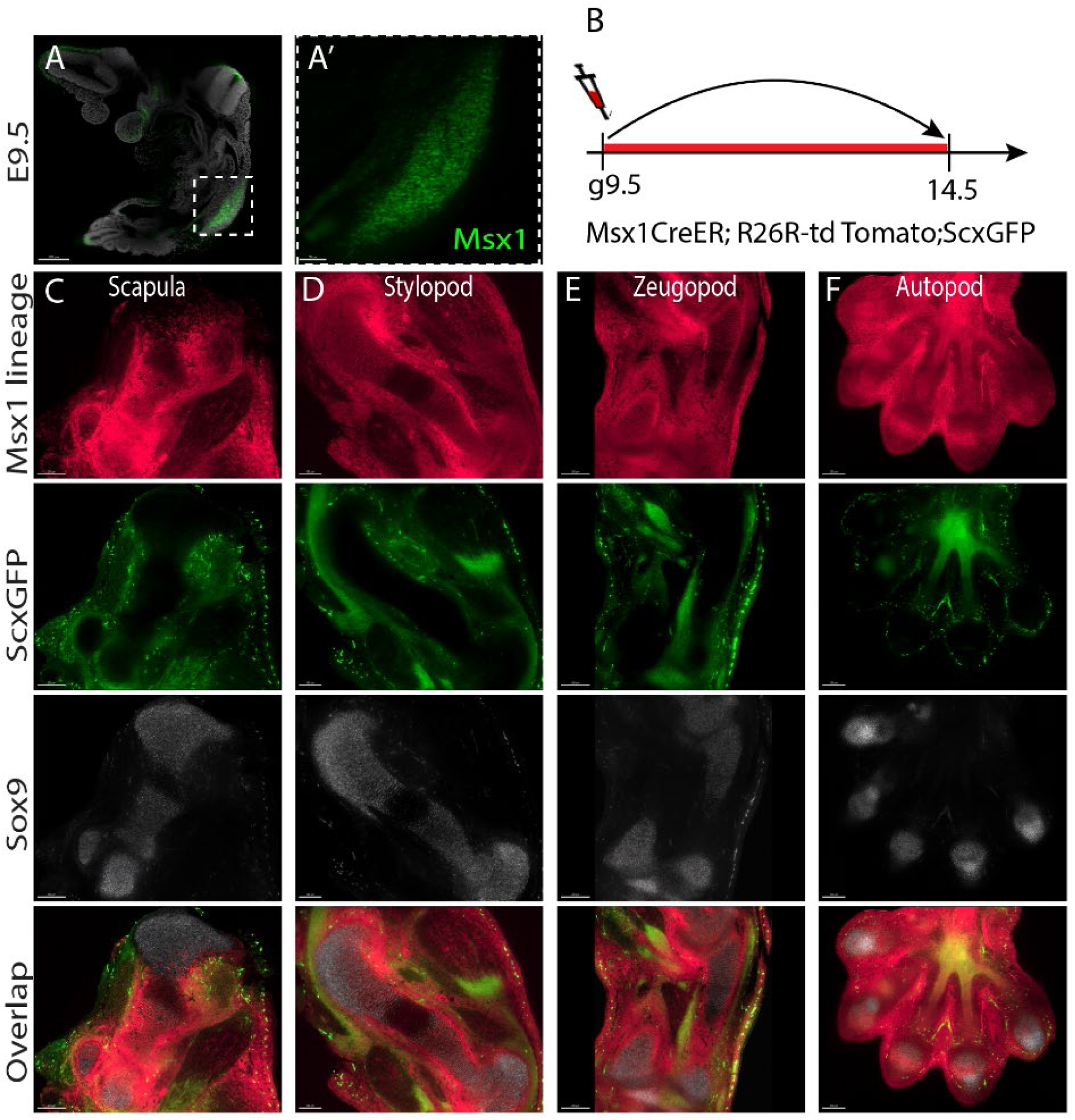
*Msx1* marks the naïve progenitors of the limb. (A) Optical section through E9.5 embryo stained for *Msx1* mRNA (green) using *in situ* HCR, counterstained with DAPI (grey) and imaged by light sheet microscopy. Dashed white square demarcates the forelimb. (A’) Magnification of dashed white square in A shows that at E9.5, *Msx1* is expressed throughout the forelimb mesenchyme (n=2). (B) Scheme showing the design of the pulse-chase cell lineage experiment. *Msx1*^+^ cells were marked at E9.5 by administration of a single dose of tamoxifen to *Msx1-CreER^T2^; Rosa26-tdTomato, Scx-GFP* pregnant females (g, gavage). The contribution of *Msx1* lineage to the limb tissues was examined at E14.5. (C-F) Optical sections through scapula (C), stylopod (D), zeugopod (E) and autopod (F) show that at E14.5, descendants of E9.5 *Msx1^+^* cells (pink) contribute to all mesenchyme-derived tissues of the forelimb and most of the scapula, including tendons (green, visualized by *Scx-GFP*), cartilage (grey, visualized by *in situ* HCR staining of *Sox9* mRNA), muscle connective tissue and perichondrium (n=3). Whole forelimbs were imaged by light sheet microscopy.

Our single-cell data suggested that during developmental, *Msx1* naïve progenitors progressively transition into proximal and autopodial progenitors. To demonstrate this process *in vivo*, we combined sequential pulse-chase genetic lineage tracing, using the *Msx1-CreER^T2^; Rosa26-tdTomato* mice, with whole-mount *in situ* HCR for *Msx1*. By applying sequential short (30 h) chase periods, we could follow the temporal dynamics of *Msx1* cell differentiation, whereas *in situ* HCR provided the position of the *Msx1* naïve progenitors at the end of the chase. Differences in position between *Msx1^+^* naïve progenitors and their descendants would provide spatiotemporal information on the progression of this process. To demonstrate the transition of *Msx1^+^* naïve progenitors to proximal progenitors we performed HCR for *Shox2*, which marks the latter cells.

At E10.5, while restricted *Msx1* expression was observed at the outer margin of the limb along the anterior-posterior axis, tdTomato signal was detected throughout the limb, including in *Msx1*^+^ cells. *Shox2* expression was observed in *tdTomato^+^Msx1^-^* cells, which were surrounded by *tdTomato^+^Msx1^+^* cells (Figure 4A). These results indicate that at that stage, some of the pulsed cells maintained their naïve state, mostly at the margin of the limb, whereas the center was occupied by *Msx1* lineage cells that had lost their naïve state and adopted a new fate, some as proximal progenitors. At E11-E11.5, *Msx1* expression in the outer margin of the limb was maintained, whereas tdTomato-expressing cells were found in the anterior proximal domain but not in the proximal posterior domain. *Shox2* expression was observed in the center of the limb, overlapping the tdTomato signal except at the posterior side. These results demonstrate the transition of *Msx1* naïve progenitors into proximal progenitors between E10.5 and E11.5 (Fig.4B-C). At E12.5, *Msx1* expression was observed in the autopod margin and interdigital space, demarcating metacarpals 2-5. tdTomato-expressing cells were found in the autopod margin and a few in the anterior proximal domain. *Shox2* expression was observed in the proximal limb, stylopod and zeugopod, where the anterior and posterior sides adjacent to the autopod overlapped partially with the tdTomato signal (Fig. 3D), demonstrating that the transition from *Msx1* naïve progenitors into proximal progenitors was still taking place between E11.5 and E12.5. Expression of *Msx1* also by autopodial P3 progenitors raised the possibility that proximal P2 cells are derived from P3 as well. However, P3 cells also expressed *Hoxa13*, whose lineage was previous shown to contribute only to the autopod (Scotti et al., 2015b), negating this possibility.

**Figure 4.**
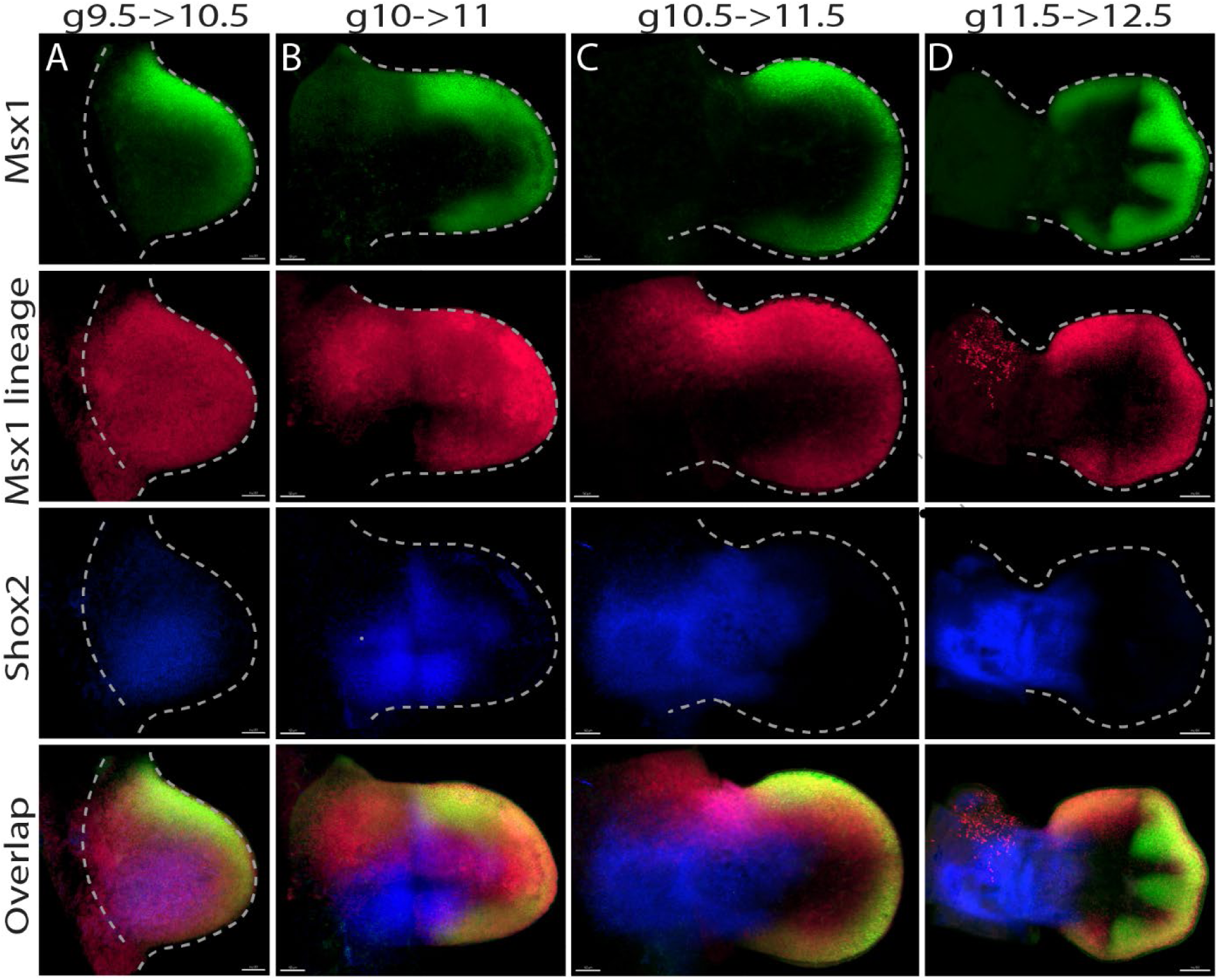
*Msx1^+^* naïve limb progenitors progressively differentiate into proximal or autopodial progenitors. (A-D) Pulse-chase experiment using *Msx1-CreER^T2^; Rosa26-tdTomato* mice. *Msx1^+^* cells were labeled by tamoxifen administration at E9.5 (A), E10.5 (B), E11 (C) or E11.5 (D) and forelimbs were harvested 30 hours later. MIP images of whole-mount forelimbs stained for *Msx1* (green) and *Shox2* (blue) mRNA using *in situ* HCR and imaged by light sheet microscopy show that descendants of *Msx1^+^* cells (pink) contribute to proximal (*Shox2^+^*) and autopodial (*Shox2^-^*) limb domains progressively and concurrently. At each stage, n=2. g, gavage.

Finally, we examined the level of cell fate stability of the *Msx1^+^* naïve progenitors during development. For that, we compared between *Msx1^+^* naïve progenitors from different time points for differentially expressed genes. As seen in Fig. S3A-F, *Igdcc3*, *Asb4*, and *Hmga2* were found to be highly expressed by E10.5 and E11.5 P1 cells as compared to later stages, whereas E12.5 P1 cells displayed higher expression of the autopodial marker *Hoxd13*. Comparison between E12.5 and E13.5 P1 cells did not reveal differentially expressed genes. Overall, this analysis indicates that the naïve progenitors undergo a mild transcriptional change over time, as they largely maintain their transcriptional program.

Together, these results confirm and expand our single-cell data and show that *Msx1^+^* progenitors are the most naïve multipotent mesenchymal progenitors, which give rise to all mesenchyme-derived tissues of the limb. We demonstrated the progressive transition of *Msx1^+^* naïve progenitors to proximal progenitors, which takes place for several days, concomitantly with autopod development. Moreover, we showed that the addition of new proximal progenitors took place mainly on the anterior side and, to a lesser extent, on the posterior side.

### Differentiation of *Msx1* lineage cells to *Sox9^+^* chondroprogenitors occurs progressively and simultaneously along the different skeletal segments

Having found progressive differentiation of *Msx1^+^* naïve progenitors, we proceeded to study the dynamics of the differentiation of this lineage into chondroprogenitors by comparing the expression of *Msx1* to that of *Sox9*, the earliest known chondro-osteogenic marker (Akiyama‡ et al., 2005; Bi et al., 1999; Lefebvre et al., 1997). For that, we established a chondrogenic gene module anchored to *Col2a1*, a *bona fide* chondrogenic marker (Benoit de CrombruggheU, Veronique Lefebvre and Weimin Bi, Shunichi Murakami, 2000), and used it to compute a chondrogenic score. Cells from each day were ordered by chondrogenic score and binned into 65 bins. The mean expression of *Sox9* and *Msx1* was calculated for each bin, and trend line and confidence interval were calculated (see Methods). This analysis revealed that at all sampling time points, cells with low chondrogenic score expressed high levels of *Msx1* and low levels of *Sox9*. As cells progressed through differentiation, *Sox9* expression was upregulated as expected, while *Msx1* expression was downregulated (Fig. 5A,B). To validate this, we performed *in situ* HCR for *Sox9* and *Msx1* on E10.5-E12.5 forelimbs. As seen in Figure S4A-C and in agreement with the single-cell results, the expression domains of *Msx1* and *Sox9* were mutually exclusive, with slight overlap at the borders, which likely represents the transitional stage.

**Figure 5.**
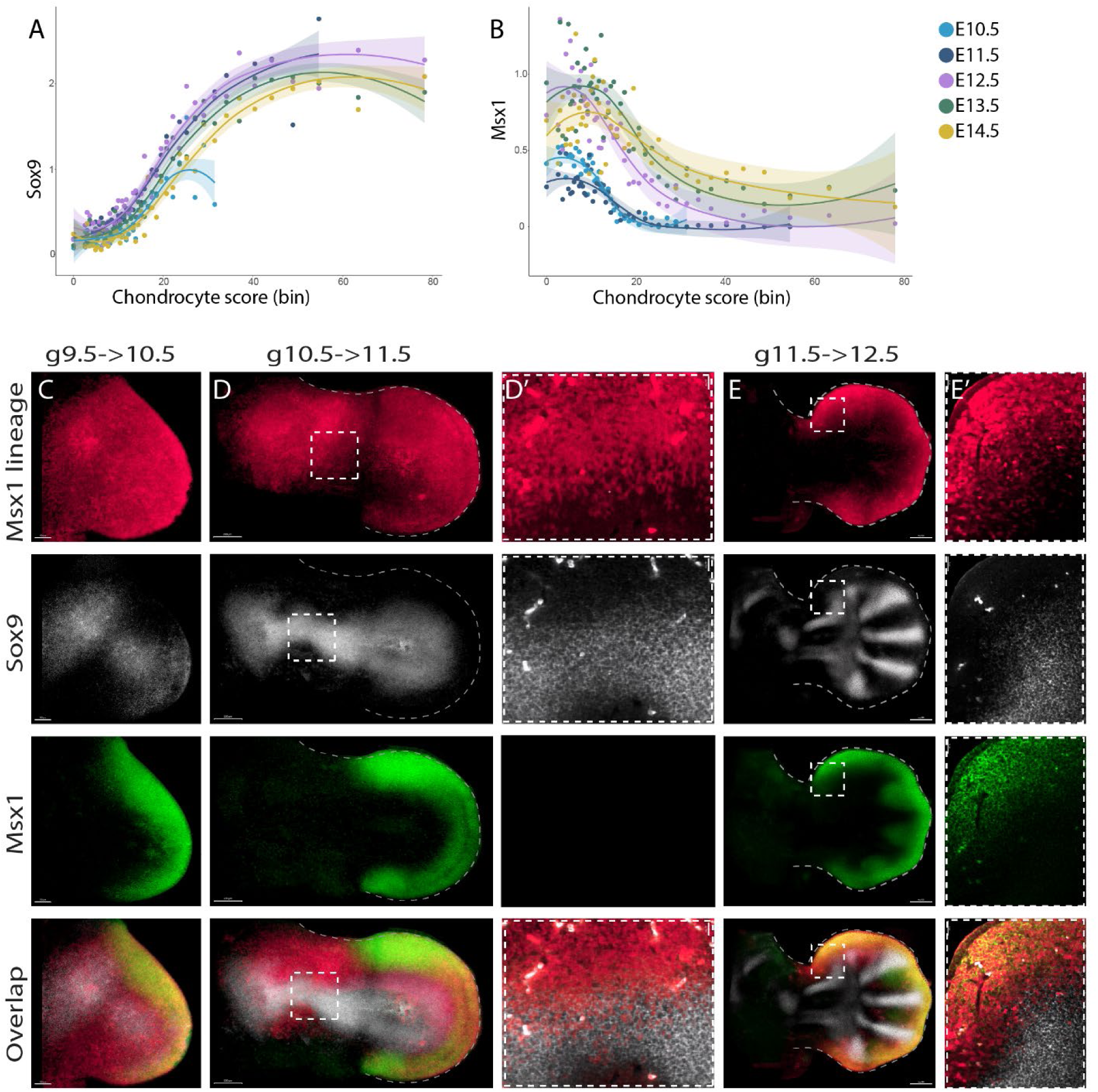
Spatiotemporal analysis of the differentiation of *Msx1^+^* progenitors into *Sox9^+^* cells. (A,B) Graphs showing the expression of *Sox9* (A) and *Msx1* (B) by progenitor cells and chondrocytes that were ordered by chondrogenic scores. As cells begin to differentiate into chondrocytes, *Msx1* is downregulated concurrently with *Sox9* upregulation. Marker genes for proximal and autopod limb segments, as well as the 10 most significantly differentially expressed genes, are indicated by red dots. (C-E) Pulse-chase experiment using *Msx1-CreER^T2^; Rosa26-tdTomato* mice. *Msx1^+^* cells were labeled by tamoxifen administration at E9.5 (C), E10.5 (D,D’) or E11.5 (E,E’) and forelimbs were harvested 30 hours later. Whole-mount forelimbs were stained for *Msx1* (green) and *Sox9* (grey) mRNA using *in situ* HCR and imaged by light sheet microscopy. MIP images show that descendants of *Msx1^+^* cells (pink) differentiate into *Msx1^-^Sox9^+^* cells progressively and simultaneously in different parts of the skeleton. D’ and E’ are magnifications of the dashed white squares in G and H, respectively. At each stage, n=2. g, gavage.

Our data analysis indicated that the transition from *Msx1* to *Sox9* expressing cells takes place in each of the examined days, suggesting that the differentiation of *Msx1* naïve progenitors into chondroprogenitors is a progressive process. To demonstrate *in vivo* the spatiotemporal dynamics of this process, we combined sequential short (30 h) pulse-chase experiments using the *Msx1-CreER^T2^; Rosa26-tdTomato* mice with whole-mount *in situ* HCR for *Msx1* and *Sox9*. At E10.5, *Msx1* lineage cells populated the entire limb. *Sox9* expression was observed in the center of the limb, within a *tdTomato^+^Msx1^-^* domain, which was surrounded by *tdTomato^+^Msx1^+^* cells at the margin (Fig. 4C). Examination at E11.5 showed that *Msx1* lineage cells populated most of the limb, excluding the most proximal posterior domain. *Sox9* expression domain was observed in the center of the limb. Interestingly, the *Sox9* expression domain overlapped with *tdTomato^+^Msx1^-^* cells along the entire forming skeleton, mostly on the anterior side (Fig. 5D,D’). *tdTomato^+^Msx1^+^* cells were located at the autopod margin, surrounding the distal *Sox9* expression domain. At E12.5, *Msx1* lineage cells were observed at the autopod margin, with a small proximal extension on the anterior side. *Sox9* expression demarcated the humerus, radius, ulna, and five metacarpals. *Sox9* expression overlapped with *tdTomato^+^Msx1^-^* cells at the distal anterior radius, metacarpals 1 and 5 and tips of metacarpals 2-4. Areas of overlap between *Sox9* expression and *tdTomato^+^Msx1^+^* cells were detected in the lateral side of metacarpals 2-4 and at all metacarpal tips (Fig. 5E,E’).

Overall, these results support the temporally progressive differentiation of *Msx1* lineage cells not only to proximal and autopodial progenitors, but also into *Sox9^+^* chondroprogenitors. Moreover, this process is not restricted spatially, but rather occurs simultaneously along all segments of the developing skeleton.

### The skeleton forms progressively and nonconsecutively in a complex three-dimensional pattern

Our observation that the progressive differentiation of *Msx1^+^* naïve progenitors to chondroprogenitors occurs simultaneously in all segments of the developing limb skeleton prompted us to reexamine the order by which these segments form. The finding that *Msx1* expression is lost once the naïve progenitors differentiate provided us with a unique opportunity to address this question. The rationale behind our approach was that the first element to form would be composed of descendants of progenitors that lost *Msx1* expression first, whereas the last element to form would be composed of descendants of progenitors that were last to lose *Msx1* expression. To follow temporally the loss of *Msx1* expression by naïve progenitors, we performed consecutive pulse-chase lineage tracing experiments by administering single doses of tamoxifen to *Msx1-CreER^T2^; Rosa26-tdTomato* mice at E9.5, E10.5, E11.5 or E12.5. To determine the exact 3D spatial distribution of tdTomato-positive cells in the forming skeleton, we cleared limbs of E13.5 embryos and imaged them using light sheet microscopy.

As seen in Figure 6A, tamoxifen administration at E9.5 resulted in tdTomato labeling of the entire skeleton. However, pulsing at E10.5 (Fig. 6B) resulted in loss of tdTomato signal in most of the scapula, ventral humerus (Fig. S5A) and radius. tdTomato signal was observed in the acromion, humeral head and deltoid tuberosity, dorsal humerus, ulna, and in the entire autopod (Fig. 6B, Fig. S5B). Pulsing at E11.5 (Fig. 6C) resulted in loss of tdTomato signal in acromion and humeral shaft and metacarpals of digits 3 and 4 (Fig. S5C). Still, tdTomato signal was detected at the humeral head and deltoid tuberosity, radius and most of the digits. Finally, following pulsing at E12.5 (Fig. 6D), tdTomato signal was lost in the radius and metacarpals, but remained at the tips of the growing digits (Fig. S5D). These results indicated that the skeleton forms progressively in a complex pattern and not linearly along the proximal-distal axis.

**Figure 6.**
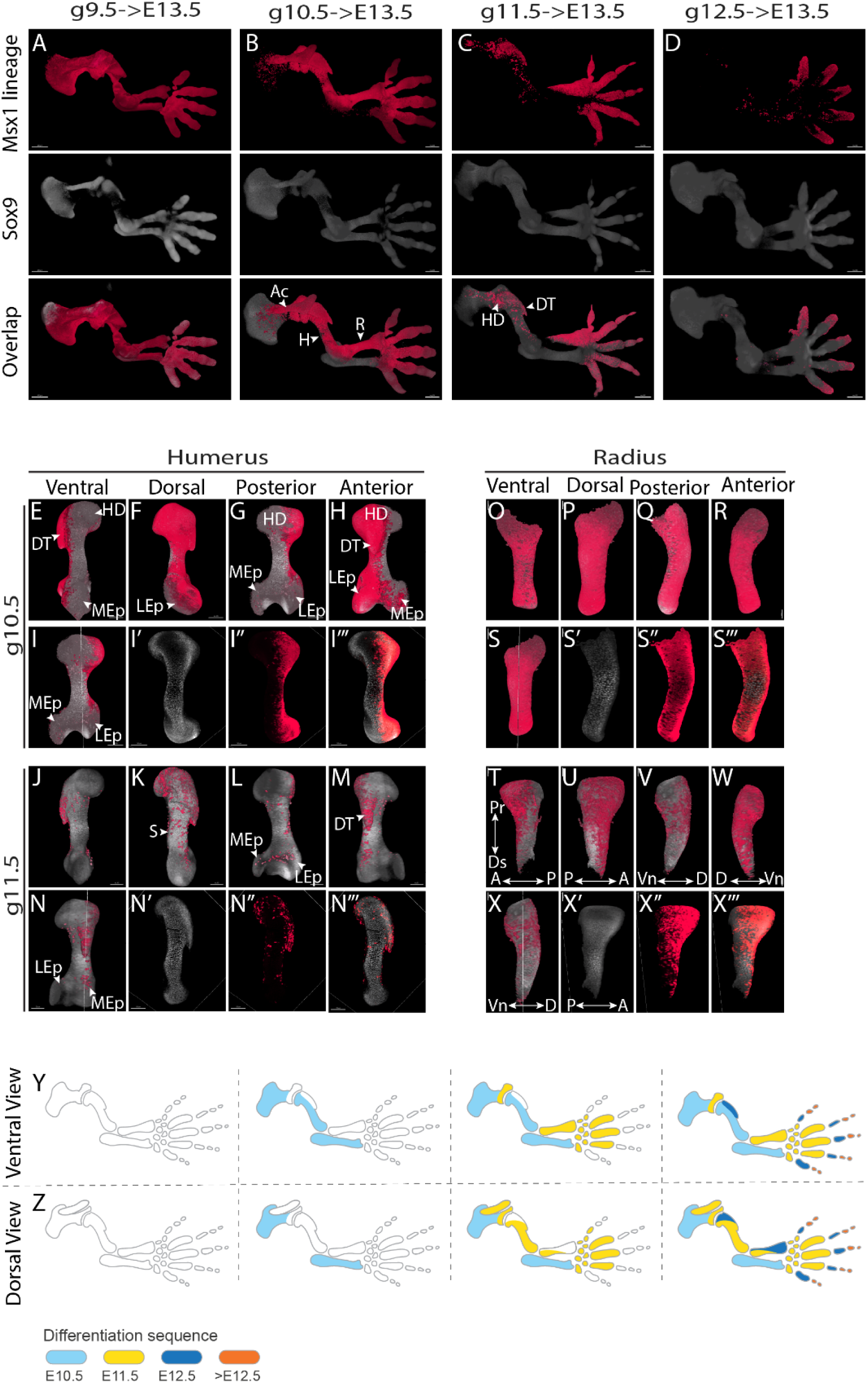
*Msx1* lineage tracing reveals that the patterning of skeletal element deviates from the PD model. (A-X) *Msx1^+^* cells were marked at consecutive days from E9.5 to E12.5 by administration of single doses of tamoxifen to *Msx1-CreER^T2^; Rosa26-tdTomato* pregnant females. The contribution of *Msx1* lineage cells to the limb skeleton was examined at E13.5. Whole-mount forelimbs stained with anti-SOX9 antibody (grey) and imaged by light sheet microcopy show that *Msx1* lineage (pink) contributes to skeletal formation in a complex, non-linear pattern. (A-D) Dorsal view of 3D-rendered images of pulse-chase cell lineage experiment. Descendants of *Msx1^+^* cells marked at E9.5 (A) were detected throughout the skeleton (n=4), whereas pulsing at E10.5 (B) resulted in tdTomato signal in acromion, humerus, radius and in the autopod (n=4). Following pulsing at E11.5 (C), tdTomato signal was detected in humeral head, deltoid tuberosity, part of the radius and digits (n=3). Finally, pulsing at E12.5 (D) labeled the tips of the digits (n=2). (E,J) Ventral, (F,K) dorsal, (G,L) posterior and (H,M) anterior views of 3D-rendered humerus images following pulsing at E10.5 (E-I’’’) and E11.5 (J-N’’’) show that the humeral ventral side forms first, followed by dorsal epicondyle and shaft, whereas the humeral head and deltoid tuberosity are the last to form. The locations of optical sections shown in I’-I’’’ and N’-N’’’ are indicated in I,N. (O,T) Ventral, (P,U) dorsal, (Q,V) posterior and (R,W) anterior views of 3D-rendered radius images following pulsing at E10.5 (O-S’’’) and E11.5 (T-X’’’) show that the ventral side of the radius forms first, followed by formation of the dorsal side in a diagonal AP direction. The locations of optical sections shown in S’-S’’’ and X’-X’’’ are indicated in S,X. (Y-Z) Schematics showing the spatiotemporal differentiation sequence in the forelimb skeleton from a ventral (Y) and dorsal view (Z). Abbreviations: Ac, acromion; R, radius; H, humerus; DT, deltoid tuberosity; HD, humeral head; LEp, lateral epicondyle; MEp, medial epicondyle; S, shaft; Pr, proximal; Ds, distal; A, anterior; P, posterior; Vn, ventral; D, dorsal; g, gavage.

To better understand this process, we examined in greater detail the tdTomato signal in the humerus and radius. As seen in Figures 6E,M’’’, in the humerus, pulsing at E10.5 led to the loss of tdTomato signal at the ventral side from head to medial epicondyle, whereas the dorsal side was tdTomato-positive from head to lateral epicondyle. Following pulsing at E11.5, the areas of the humerus that lost tdTomato signal were in the dorsal shaft and lateral epicondyle, whereas the deltoid tuberosity and the dorsal side of humeral head were still tdTomato-positive (Fig. 6J-N’’’).

In the radius, tamoxifen administration at E10.5 resulted in tdTomato labeling throughout the bone (Fig. 6O-S’’’). However, pulsing at E11.5 resulted in loss of tdTomato signal in almost the entire ventral side, with few labeled cells on its distal tip (Fig. 5T). On the dorsal side, the proximal posterior side of the radius was mostly tdTomato-negative, while the anterior side still displayed extensive labeling (Fig. 6U-X’’’). These results indicate that the ventral side of the radius forms first, followed by a diagonal AP direction of dorsal radius formation. These results demonstrate that the skeleton form in a complex pattern that extends to the level of the single element.

To validate the results of the *Msx1* lineage experiments, we examined the spatiotemporal dynamics of chondroprogenitor differentiation in the limb by following the induction of *Sox9* expression. For that, we used the *Sox9-CreER^T2^* mice, which were previously shown to efficiently drive the expression of *Rosa26-lacZ* reporter in the developing skeleton (Soeda et al., 2010). Because both our single-cell analysis and lineage studies showed that at E10.5, only part of the *Sox9* chondroprogenitors were differentiated (Fig. 6B), we decided to activate Cre activity at this time point. Thus, we administered a single dose of tamoxifen at E10.5 and harvested the limbs 30 h and 72 h afterwards. E11.5 whole-limbs were stained for *Sox9* mRNA using *in situ* HCR, whereas E13.5 whole limbs were stained for SOX9 protein and imaged using light sheet microscopy. As shown in Figure 7A-E’’’, tdTomato signal was observed in most of the scapula, and only on the ventral-posterior side of the humerus and ulna, whereas the radius and autopod were tdTomato-negative. At 72 h post-induction (Fig.67F-J’’), tdTomato labeling was seen in most of the scapula, the entire humeral shaft and ulna; however, the acromion, humeral head, deltoid tuberosity, lateral and medial epicondyles, elbow and most of the radius and digits were tdTomato-negative. These results further support the data obtained using the *Msx1-CreER^T2^* line, demonstrating that the skeleton forms nonconsecutively in a complex pattern that involves not only the PD axis, but also the DV and AP axes, extending to the level of the single element.

**Figure 7.**
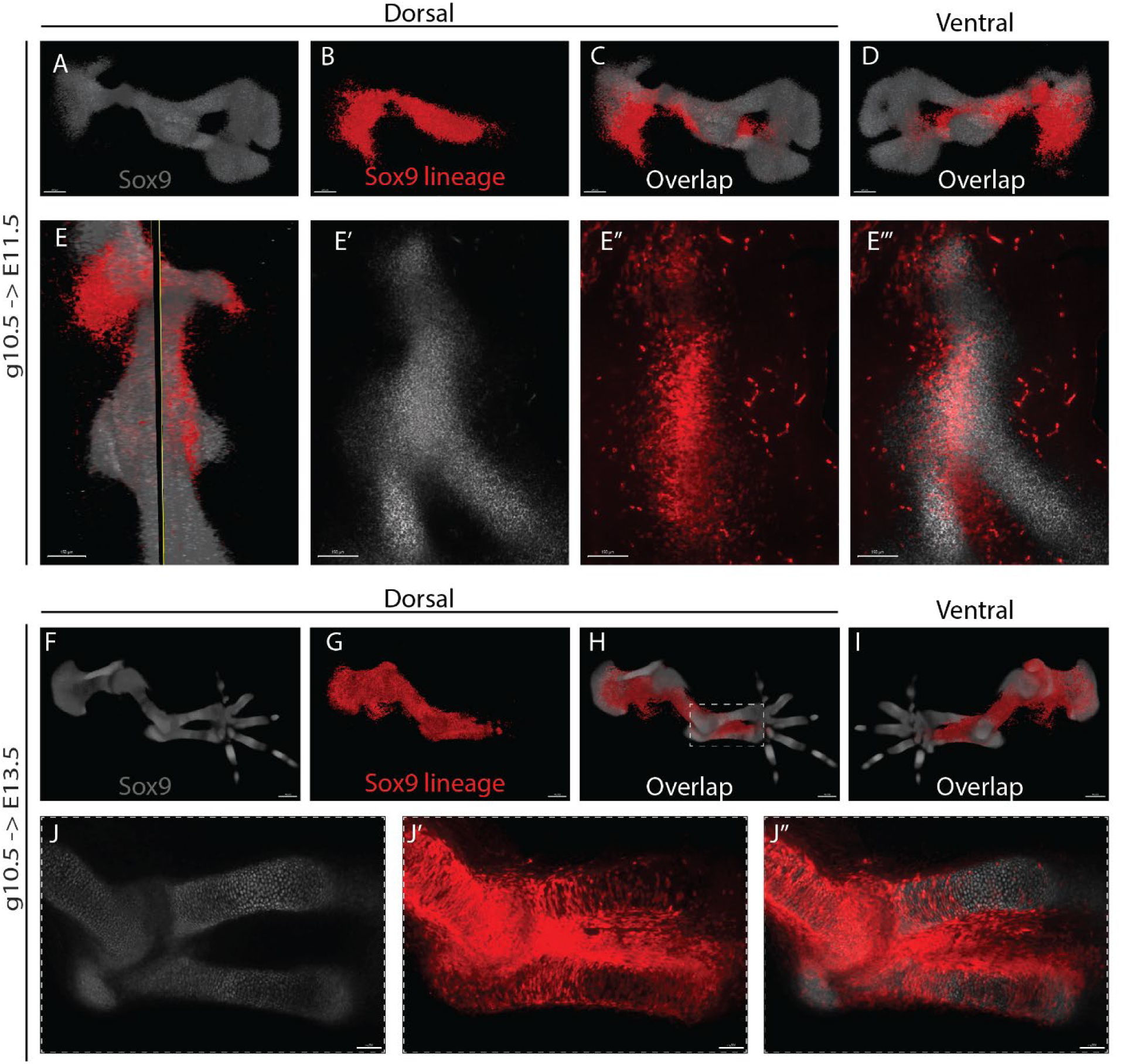
*Sox9* lineage tracing confirms that skeletal chondroprogenitor differentiation occurs progressively in a complex 3D pattern. Pulse-chase experiment using *Sox9-CreER^T2^; Rosa26-tdTomato* mice. *Sox9^+^* cells were labeled by tamoxifen administration at E10.5 and forelimbs were harvested 30 hours later. Forelimbs were stained for *Sox9* (grey) mRNA using *in situ* HCR and imaged by light sheet microscopy. (A-D) 3D-rendered images show that pulsing of *Sox9*^+^ cells at E10.5 results in incomplete labeling of the skeleton, demonstrating the progressive differentiation into *Sox9^+^* cells in a complex, non-consecutive pattern. The locations of optical sections shown in E’-E’’’ is indicated in E. (F-I) 3D-rendered images of *Sox9-CreER^T2^; Rosa26-tdTomato* mice forelimbs labeled by tamoxifen administration at E10.5 and harvested 72 hours later. Forelimbs were stained for SOX9 protein (grey) and imaged by light sheet microscopy. Labeling of *Sox9*+ cells at E10.5 results in incomplete labeling of the skeleton. (J-J’’) Optical section through zeugopod segment (demarcated by dashed white square in H) shows that cells of the ulna upregulate *Sox9* expression prior to cells of the radius.

## DISCUSSION

In this work, we revisit the long-standing question of the spatiotemporal sequence of limb development using modern molecular tools. We generated a comprehensive cellular atlas of the limb mesenchymal cell lineages during development. Using this atlas, we identified a population of naïve progenitors and their progressive and simultaneous transition into proximal and autopodial progenitors. We establish *Msx1* as a marker of naïve progenitors and localize them to the outer margin of the developing limb, along the anterior-posterior axis. We then showed that the descendants of these progenitors progressively contribute to the entire forming skeleton. Finally, temporal analysis of the differentiation of naïve progenitors revealed that the skeleton forms progressively in a complex 3D pattern, which extends to the single element level (Fig. 8).

**Figure 8.**
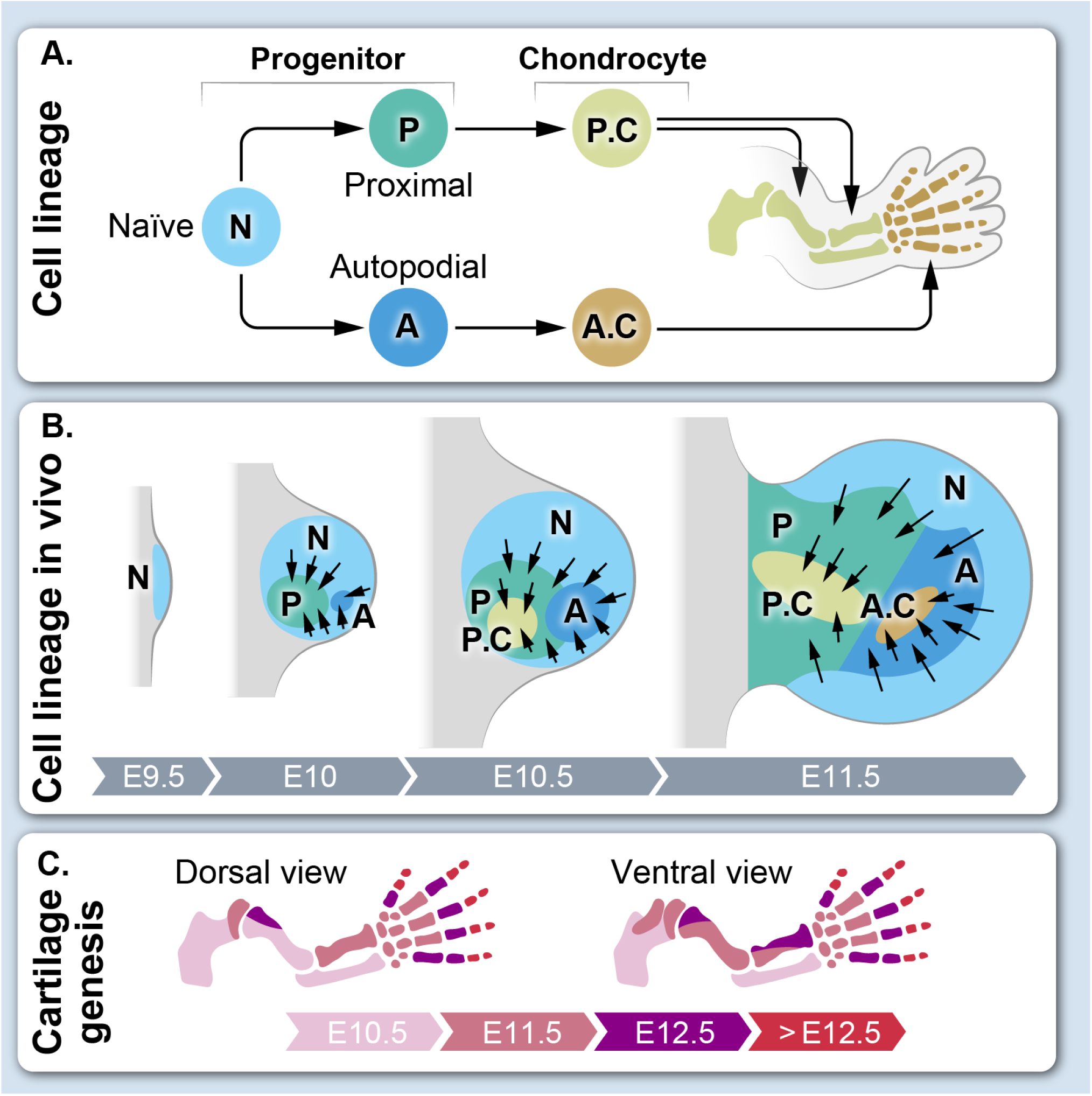
Model for the patterning and development of the limb skeleton A. *Msx1^+^* cells are naïve, multipotent limb mesenchymal progenitors that give rise to proximal and autopodial progenitors that, in turn, differentiate into proximal and autopodial chondrocytes, respectively. **B.** In the developing limb, *Msx1^+^* naïve progenitors undergo specification first into proximal and, soon after, into autopodial progenitors. This progressive process continues for several days. Concurrently, already specified proximal and autopodial progenitors start to differentiate into chondroprogenitors by upregulating *Sox9* expression. This process occurs simultaneously at different locations along the forming limb. **C.** A new model for skeleton development. The skeleton forms progressively and simultaneously from multiple foci, in a nonconsecutive fashion along the proximal-distal, dorsal-ventral and anterior-posterior axes. For example, proximal scapula, dorsal humerus and ulna form together, followed by more distal scapula, ventral humerus, proximal radius and middle metacarpals.

Our systematic single-cell analysis of limb mesenchymal cells revealed a pool of naïve progenitors. These cells transit initially to proximal progenitors and, shortly afterwards, also to autopodial progenitors. The finding that the pool of *Msx1* naïve progenitors is maintained for several days suggests that this transition occurs progressively. These finding correspond with some aspects of previously suggested models of limb development. The existence of naïve limb progenitors and their progressive transition into progenitors of the different limb segments was suggested by the progress zone model (Saunders, 1948; Summerbell et al., 1973; Wolpert, 2002). The coexistence of the different progenitors of these segments was suggested by the early specification model, whereas the progressive and concurrent specification of proximal and distal fates is consistent with the two-signal model (Mariani et al., 2008; Mercader et al., 1999a, 2000). In this respect, our findings integrate the three models.

Because the limb skeleton comprises three segments, it was expected to originate from three different pools of progenitors. However, in line with previous studies that failed to identify zeugopod-specific markers (Tabin and Wolpert, 2007b), we also were unable to subdivide the proximal gene program into stylopod- and zeugopod-specific. This finding raises the question of the mechanism underlying the formation of the stylopod and zeugopod as two separate segments.

Recently, several works studying limb development at single-cell resolution have been published (Desanlis et al., 2020a; Feregrino et al., 2019; Kelly et al., 2020). Our work is unique in that we sampled cell from the entire limb daily from E10.5 up to E14.5, providing a complete and continuous representation of cell states and transcriptional changes that take place in the limb during this critical stage of limb patterning and differentiation.

Our single-cell data revealed a set of markers for the three progenitor populations we identified. While *Msx1, Lhx9,* and *Lhx2* marked the naïve progenitors, *Msx1* lineage cells gave rise to all mesenchyme-derived limb tissues. These TFs are involved in FGF, BMP and Shh signaling, major pathways that regulate patterning along the PD, AP and DV axes (ALAPPAT et al., 2003; Bensoussan-Trigano et al., 2011; Lallemand et al., 2005, 2009; Tzchori et al., 2009; Watson et al., 2018; Yang and Wilson, 2015). Interestingly, we found that the naïve progenitors largely maintain their transcriptional program during limb development. For the proximal progenitors, we identified a set of markers that includes *Meis1* and *Meis2*, two well-known proximal markers (Capdevila et al., 1999; Delgado et al., 2020; Mercader et al., 1999b, 2009), *Shox2, Pkdcc* and many other genes. The validity of *Shox2* as a marker for proximal cells is supported by lineage studies showing that *Shox2* lineage gives rise to the proximal part of the limb, ending at the wrist (Sun et al., 2013). For the autopod progenitors, we identified a set of markers that included *Hoxd13*, *Hoxa13* and *Hoxd12*. These genes are expressed specifically in the autopod and play an essential role in digit identity and patterning (Desanlis et al., 2020b; Fabre et al., 2018; Fromental-Ramain et al., 1996; Knezevic et al., 1997; Scotti et al., 2015a; Sheth et al., 2007; Zákány et al., 1997). These markers were co-expressed with *Msx1*, *Msx2*, *Lhx9*, and *Lhx2*, which we show to be important for autopod patterning, suggesting a functional link between these two groups of genes.

The identification of markers for the three progenitor populations allowed us to study their spatiotemporal distribution during limb development. As suggested by the single-cell analysis results, we found that the naïve progenitors marker *Msx1* was expressed throughout the process in an arc-like pattern along the anterior-posterior axis, as well as dorsally and ventrally away from the AER. This finding suggests that the naïve progenitors maintain their location through development. Moreover, it suggests that their progressive transition to the different lineages may not be restricted to the limb apex, but can occur along the anterior-posterior axis. Indeed, sequential pulse-chase experiments clearly showed that between E9.5 and E11.5, *Msx1* lineage cells populated extensive areas in the proximal side of the limb. Moreover, they overlapped with both proximal and autopodial progenitors. Proximal progenitors were initially located in the center and later expanded proximally, whereas autopodial progenitors were initially located distally in the posterior-dorsal side and later expanded anteriorly. We therefore suggest that the progressive transition of naïve progenitors into proximal and autopodial fates occurs along the length of the limb, allowing simultaneous transition into these identities.

The establishment of limb progenitor identities and their differentiation to chondroprogenitors can follow two scenarios. One possibility is that the two processes are separated temporally, such that differentiation starts only after all progenitor identities have been established. Alternatively, a progressive process of identity establishment may coincide with the differentiation of the two progenitor pools into chondroprogenitors. Our findings indicate that the differentiation of naïve *Msx1^+^* progenitors into chondroprogenitors is progressive and that the transition of these naïve progenitors into proximal and autopodial progenitors coincides with the differentiation of these two progenitor pools into chondroprogenitors, suggesting that these processes overlap temporally. Strong support for this possibility is our observation that at E10.5-E11.5, both *Msx1*, which marks the naïve progenitors, and *Sox9*, which marks chondroprogenitors, were expressed in the developing limb. Other pieces of evidence that are consistent with this scenario came from genetic lineage tracing analyses in mice. We previously showed that different skeletal elements form progressively by continuous addition of *Sox9^+^* cells (Blitz et al., 2013; Eyal et al., 2019)

In this work, we provide several pieces of evidence to support the conclusion that *Msx1* is a marker for the naïve mesenchymal progenitors. These include our single-cell analysis and lineage tracing using *Msx1-CreER^T2^* mice. This knock-in allele was previously shown to drive an identical pattern of Cre expression as the endogenous *Msx1* gene (Lallemand et al., 2013). The combination of a reliable marker, mouse line and the finding that *Msx1* expression by naïve mesenchymal progenitors is lost once they differentiate provided us with a unique opportunity to study the order by which the skeleton forms. If the common view is correct and the skeleton forms in a proximal-to-distal direction, then progenitors of the proximal stylopod should be first to lose *Msx1* expression and differentiate, followed by zeugopod progenitors and, lastly, by autopod progenitors. However, the pattern that we observed was much more complex and nonconsecutive, as skeletogenesis occurred simultaneously progressing from multiple foci along the limb. This finding indicates that in addition to the formation of skeletal elements along the PD axis, there is also strong contribution along the AP and DV axes. An example for the complexity of the process is our finding that the posterior half of the humerus formed first together with ulna, whereas the anterior side of the humerus formed later together with the radius. Further support for this notion is the similar results we obtained studying the order by which the skeleton forms using the spatiotemporal elevation of *Sox9* expression in chondroprogenitors.

In summary, our findings suggest a new model for limb and skeleton development. At its core is the principle that limb development involves progressive and simultaneous transition of naïve limb progenitors into either proximal or autopodial progenitors, which then progressively differentiate into *Sox9^+^* chondroprogenitors. This process occurs simultaneously at different locations along the limb, suggesting that the skeleton forms progressively from multiple foci in a complex 3D pattern.

## METHODS

### Animals

All experiments involving mice were approved by the Institutional Animal Care and Use Committee (IACUC) of the Weizmann Institute. The generation of *Sox9-CreER^T2^* (Soeda et al., 2010), *Scx-GFP* (Pryce et al., 2007), *Msx1-CreER^T2^* (Lallemand et al., 2013), *Sox9-GFP* (Chan et al., 2011) and *Rosa26-tdTomato* (Madisen et al., 2010) mice has been described previously. For fluorescence-activated cell sorting (FACS) experiments, *Sox9-CreER^T2^-tdTomato*;*Scx-GFP* or *Sox9-GFP* mice were crossed with *Rosa26-tdTomato;Scx-GFP* or C57BL/6 mice, respectively. For lineage tracing experiments, *Msx1-CreER^T2^* were crossed with *Rosa26-tdTomato* or with *Rosa26-tdTomato;Scx-GFP* reporter mice. Plug date was defined as E0.5.

Induction of Cre recombinase was performed at indicated pregnancy stages by administration of 5 mg/ml tamoxifen in corn oil X5 body weight by oral gavage. For harvesting of embryos, timed-pregnant females were euthanized by cervical dislocation.

### Cell isolation and flow cytometry

Single-cell experiments were performed on forelimbs from E10.5, E11.5, E12.5, E13.5 and E14.5 mouse embryos. For collection of E10.5 cells, *Sox9-GFP* and *Sox9-CreER^T2^-tdTomato;ScxGFP* (without tamoxifen induction) mice were used. For collection of E11.5-E14.5 cells, *Sox9-CreER^T2^-tdTomato;ScxGFP* mice were used 48 h after Cre induction. Forelimbs were dissected and minced in cold PBS using small scissors. For each biological replicate, forelimbs of embryos from the same litter were pooled together (six forelimbs at E10.5 and E11.5, four forelimbs at E12.5 and E13.5, and two forelimbs from one E14.5 embryo). Forelimb tissues were disassociated using enzymatic digestion. E10.5-E11.5 forelimbs were digested with pre-heated 0.25% trypsin in DMEM medium (ThermoFisher) and incubated for 10 min at 37°C, gently pipetting every 3 min. E12.5-E14.5 forelimbs were digested with 1.5 mg/ml collagenase type V (Sigma-Aldrich) in DMEM at 37°C for 10-15 min, gently pipetting every 5 min until the tissue completely dissolved. The digestion reaction was stopped by addition of DMEM supplemented with 10% FBS and 1% Pen-Strep. Cell suspensions were filtered through a 40-µm nylon mesh and collected by centrifugation at 1000 rpm for 7 min at 4°C. Supernatant was removed and cells were resuspended in 500 µl ice-cold MACS buffer (with 0.5% BSA and 2 mM EDTA in PBS) and used immediately for FACS.

Flow cytometry analysis and sorting were performed using an AriaFusion instrument (BD Biosciences, San Jose, CA) equipped with 488, 407, 561 and 633 nm lasers, using a 100-µm nozzle. Sorting gates and fluorescence compensation were defined based on GFP, tdTomato single-stained and unstained control cells. Live cells were gated using DAPI staining (1 μg/ml) and by size and granularity using FSC-A versus SSC-A. FSC-W versus FSC-A was used to further distinguish single cells. Unstained, GFP-stained only and tdTomato-stained only cells were mixed in various combinations to verify that the analysis excluded false-positive doublets. GFP was detected by excitation at 488 nm and collection of emission using 502 longpass (LP) and 530/30 bandpass (BP) filters. tdTomato was detected by excitation at 561 nm and collection of emission using a 582/15 BP filter. DAPI was detected by excitation at 407 nm and collection of emission using a 450/40 BP filter. Data were collected and analyzed using BD FACSDiva software v8.0.1 (BD Biosciences). For single-cell RNA-seq (Jaitin et al., 2014), live cells were sorted into 384-well cell capture plates containing 2 μL of lysis solution and barcoded poly(T) reverse-transcription primers. In each plate, four empty wells were used as a control. Immediately after sorting, each plate was spun down to ensure cell immersion into the lysis solution, snap frozen on dry ice and stored at −80°C until processed.

### Massively parallel single-cell RNA sequencing (MARS-Seq)

FACS-sorted cells were used for single-cell library preparation according to MARS-seq protocol, as described in (Jaitin et al., 2014). Briefly, mRNA from cells sorted into capture plates was barcoded, converted into cDNA and pooled using an automated pipeline. The pooled sample was then linearly amplified by T7 *in vitro* transcription and the resulting RNA was fragmented and converted into sequencing-ready library by tagging the samples with pool barcodes and Illumina sequences during ligation, reverse transcription and PCR. Each pool of cells was tested for library quality and concentration was assessed as described in (Jaitin et al., 2014).

### Low-level processing and filtering

RNA-seq libraries were sequenced by Illumina NextSeq500 at a median sequencing depth of 58,585 reads per single cell. Sequences were mapped to mouse reference genome (mm10), demultiplexed, and filtered as previously described (Jaitin et al., 2014), with the following adaptations. Mapping of reads was done using HISAT version 0.1.6 (Kim et al., 2015) and reads with multiple mapping positions were excluded. Reads were associated with genes if they were mapped to an exon defined by a reference set obtained from the UCSC genome browser extended by up to 2 kb for complete 3’ peak acquire. Noise level was estimated statistically on empty MARS-seq wells; median estimated noise over all experiments was 2%. Cells with less than 600 UMIs were discarded from the analysis. After filtering, cells contained a median of 2,800 unique molecules per cell. All downstream analysis was performed in R.

### Metacells modeling

We used the metacells pipeline (Baran et al., 2019) with the following specific parameters (complete script reproducing all analyses from raw data is available in GEO: GSE185940). We removed mitochondrial genes, genes linked with poorly supported transcriptional models (annotated with the prefix “RP-“) and cell cycle genes, which were identified by correlation coefficient of at least 0.1 for one of the anchor genes *Mki67*, *Hist1h1d*, *Pcna*, *Smc4*, or *Mcm3*. We then filtered cells with total fraction of mitochondrial gene expression exceeding 30% and cells with high (> 64) expression of hemoglobin genes (*Hba-a2*, *Hba-a1*, *Hbb-b2*, *Hba-x*, *Hbb-b1*). Gene features were selected for analysis using the parameter Tvm=0.2.

The gene selection strategy produced 425 marker gene features for the computation of the metacells balanced similarity graph. We used K = 150, 500 bootstrap iterations and otherwise standard parameters (500 iterations; resampling 70% of the cells in each iteration, and clustering the co-cluster matrix with minimal cluster size set to 20). We applied outlier filtering.

The resulting metacells model was annotated using the metacells confusion matrix and analysis of known marker genes. Muscle, epidermal, Schwann and immune metacells were excluded from further analysis. Next, we applied again the metacells pipeline on the remaining cells with the above-mentioned parameters.

To annotate the resulting metacells into cell types, we used the metric FP_gene,mc_, which signifies for each gene and metacells the fold change between the geometric mean of this gene within the metacells and the median geometric mean across all metacells, thus highlighting for each metacells genes that are highly overexpressed as compared to the background. Finally, we hierarchically clustered the FP most significantly changing gene table along with a set of known marker genes to identify the major cell populations.

### Defining progenitor gene module signatures and scores

To define the gene signatures of progenitor cells, we first identified modules of co-expressed genes by Pearson’s correlation across the metacells log_2_ FP_gene,mc_ expression of the 175 most variable genes. The signature genes for each progenitor state were defined by module scores. Scores were calculated for each metacell by averaging the log enrichment scores (lfp values) of the genes in the module. This approach limited the contribution of highly expressed genes to the score.

### Calculation of a chondrogenic score

To define a chondrogenic gene signature, we identified a list of genes correlated with *Col2a1* using linear correlation over metacells log enrichment scores. To avoid over-fitting of the modeling, TFs were excluded from the list. We used this chondrogenic signature for calculation of chondrogenic score across all cells. Chondrogenic scores were calculated for each cell by averaging the metacells log enrichment scores (lfp values) of the genes in the signature. Finally, to study gene expression during chondrogenic differentiation, cells were arranged by increasing ranges of signature scores and binned into 65 bins. To validate our approach, we tested differential expression of TFs that were not part of the signature gene set and are upregulated during chondrogenesis.

### Statistical Analyses

Differential gene expression analysis was performed on log_2_ sum of UMIs normalized by reads per cell, divided by cell number. P-values were calculated using Wilcoxon test to compare between mean expressions of metacells (for Fig. 2) or cells (for Figs. 3,4).

### PACT clearing

For sample preparation, E9.5-E14.5 embryos were harvested from Bl6, *Msx1-CreER; Rosa26-tdTomato, Scx-GFP* and *Msx1-CreER^T2^; Rosa26-tdTomato* timed-pregnant females and fixed in ice-cold 4% PFA in 1x PBS overnight. PFA-fixed embryos were dissected and forelimbs were cleared using PACT method (Treweek et al., 2015; Yang et al., 2014). Briefly, samples were washed in PBS, then incubated in hydrogel solution containing 4% (wt/vol) acrylamide in 1x PBS with 0.25% thermal initiator 2,2’- azobis[2-(2-imidazolin-2-yl)propane]dihydrochloride (Wako, cat. no. VA-044) at 4°C overnight. The next day, hydrogel was polymerized at 37°C for 3 hours. The samples were removed from the hydrogel, washed in PBS, and moved to 10% SDS with 0.01% sodium azide, shaking (60 rpm) at 37°C for 1-5 days, changing the SDS solution each day. Cleared samples were washed three times for 5 min with 1× PBST (PBS + 0.1% Triton X-100 + 0.01% sodium azide) at room temperature and then subjected to whole-mount *in situ* HCR or whole-mount SOX9 immunostaining.

### Whole-mount immunostaining

To detect SOX9, samples were first incubated with proteinase K (Millipore Sigma, P9290) for 10 min at room temperature, washed and post-fixed again in 4% PFA. Then, samples were incubated with 5% goat serum, 1% BSA dissolved in PBST at 4°C overnight in order to block non-specific binding of immunoglobulin. Next, samples were incubated with primary anti-SOX9 antibodies (1:100, AB5535 Millipore Sigma) in 5% goat serum, 1% BSA dissolved in PBST shaking at 37°C for 5 days. Samples were washed four times for 2 h with 1× PBST at room temperature. Next, samples were incubated with secondary Cy5 antibodies (1:100, 715-165-150, Jackson ImmunoResearch) and 1:100 DAPI (1 mg/ml) in 5% goat serum, 1% BSA dissolved in PBST shaking at 37°C for 2 days. Samples were washed four times for 2 h with 1× PBST at room temperature and then prepared for light sheet imaging. To bring the refractive index (RI) of the sample to 1.45, it was submersed in a refractive index matching solution (RIMS) prepared by dissolving 35 g of Histodenz (Millipore Sigma, D2158) in 30 ml 0.02 M phosphate buffer, shaking gently at room temperature for 1-2 days. Finally, samples were embedded in 1% low gelling Agarose (Millipore Sigma, A9414) in PBS in a glass capillary, submerged in RIMS and stored protected from light at room temperature until imaging.

### Whole-mount *in situ* hybridization chain reaction (HCR)

The *Msx1* (NM_010835.2)*, Shox2* (NM_001302358.1)*, Hoxd13* (NM_008275.4) and *Sox9* (NM_011448.4) probes and DNA HCR amplifiers, hybridization, wash and amplification buffers were purchased from Molecular Instruments. *In situ* HCR v3.0 was performed using the protocols detailed in www.molecularinstruments.com. Briefly, PACT-cleared samples were pre-incubated with hybridization buffer and incubated overnight at 37⁰C, 60 rpm with probe solution containing 1 µL of each probe in 250 µL of pre-heated probe hybridization buffer. The next day, probes were washed four times for 15 min at 60 rpm with pre-heated wash buffer, followed by two 5-min washes at room temperature with 5xSSCT. Next, samples were pre-amplified with 250 µL of amplification buffer for 5 min at room temperature and incubated with 250 µL of hairpin mixture (5 µL of hairpin h1 and hairpin h2 from 3 µM stock for each probe) overnight in the dark at room temperature. The following day, samples were washed with 5xSSCT two times for 5 min, two times for 30 min and once for 5 min at room temperature, gently shaking. For nuclear staining, samples were incubated with 1:100 DAPI/PBS solution (DAPI stock, 1mg/ml) overnight at 4⁰C, gently shaking. Finally, samples were washed twice with 2XSSC for 5 min at room temperature gently shaking and prepared for light sheet imaging as described above for SOX9-immunostained samples.

### Light-sheet fluorescence microscopy

Samples were imaged with a Zeiss Lightsheet Z.1 microscope. For each limb, a low-resolution image of the entire limb was taken with the 20× Clarity lens at a zoom of 0.36. Light-sheet fusion of images was done if necessary in Zen software (Zeiss). Tile stitching and 3D image reconstruction were performed using Imaris software (Bitplane).

## Acknowledgements

We thank Nitzan Konstantin for expert editorial assistance, Neria Sharabi from the Department of Veterinary Resources, Weizmann Institute for his help with mouse maintenance, Dr. Yoseph Addadi and Ofra Golani from the MICC Cell Observatory unit at Life Sciences Core Facilities, Weizmann Institute, for their guidance and assistance with light sheet imaging experiments and analyzes, and Dr. Tomer Meir Salame from the flow cytometry unit at Life Sciences Core Facilities, Weizmann Institute, for his assistance with FACS. We thank Efrat Davidson from the Weizmann Institute Department of Design, Photography and Printing for designing the graphic model. Special thanks to all members of the Zelzer and Amit laboratories for encouragement and advice. I.A. is an Eden and Steven Romick Professorial Chair, supported by the Merck KGaA, Darmstadt, Germany, the Chan Zuckerberg Initiative (CZI), the HHMI International Scholar award, the European Research Council Consolidator Grant (ERC-COG) 724471- HemTree2.0, an SCA award of the Wolfson Foundation and Family Charitable Trust, the Thompson Family Foundation, an MRA Established Investigator Award (509044), the Helen and Martin Kimmel award for innovative investigation, the NeuroMac DFG/Transregional Collaborative Research Center Grant.

This study was supported by grants from the David and Fela Shapell Family Center for Genetic Disorders and by The Estate of Mr. and Mrs. van Adelsbergen (to E.Z).

## Contributions

S.M. designed, performed, and analyzed experiments; performed flow cytometry sorting experiments, annotated and interpreted the single-cell data, performed and analyzed imaging experiments. M.Z. performed flow cytometry sorting experiments and library preparations. E.D and A.G performed single-cell bioinformatics analyses. S.M, M.Z, E.D, A.G, I.A and E.Z generated the figures. S.M, E.Z and I.A wrote the manuscript. E.Z and I.A supervised the project. All authors discussed the results and commented on the manuscript at all stages.

## Ethics declarations

### Competing interests

The authors declare no competing interests.

---

**Figure S1.**
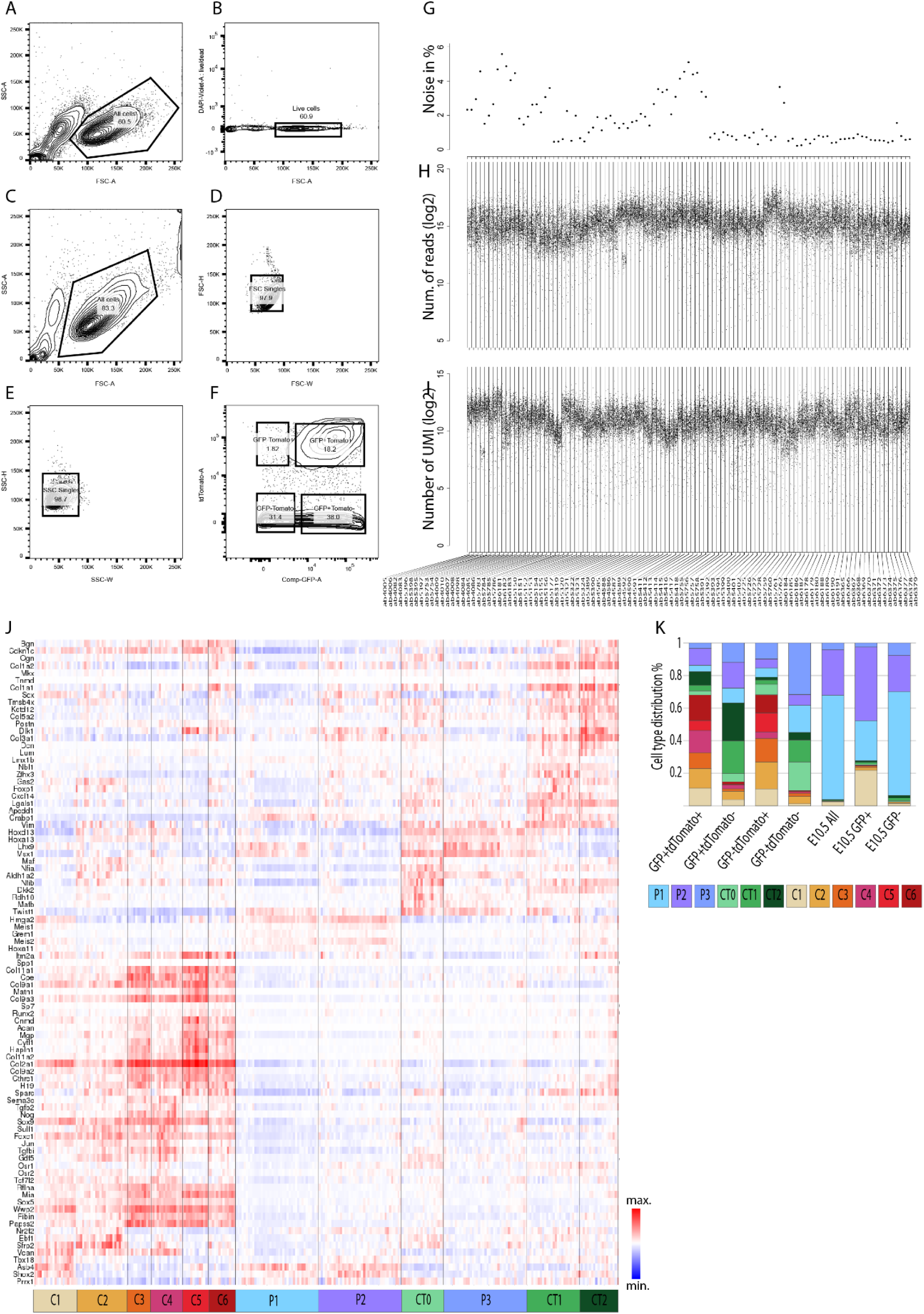
Single-cell RNA sequencing of mouse forelimb mesenchymal cells during embryonic development. Related to Figure 1. (A) Live/dead cell identification based on size (FSC-Area) and granularity (SSC-Area) gating. (B) Validation of live/dead cell gating strategy in A by DAPI staining. (C-F) Sorting strategy for isolation of single cells for MAR-seq. (C) Gating of live/dead cell based on size (FSC-A) and granularity (SSC-A). (D) Gating of single cells based on size FSC-Height versus FSC-Width. (E) Second gating for single cells based on granularity Height (SSC-H) versus Width (SSC-W). (F) Gating of tdTomato^+^; tdTomato^+^-GFP^+^; tdTomato^-^-GFP^+^; and tdTomato^-^-GFP^-^ populations. (G-I) Quality control of 32,000 analyzed single cells from the entire study. (G) Estimated ambient noise per amplification batch (103 in total). (H) Number of Illumina reads and (I) number of UMIs per amplification batch. (J) Heatmap showing log_2_ fold change in expression of key markers across metacells. Lower panels indicate association to cell types, color-coded as in Figure 1. (K) Comparison of cell type distribution between different sorting gates.

**Figure S2.**
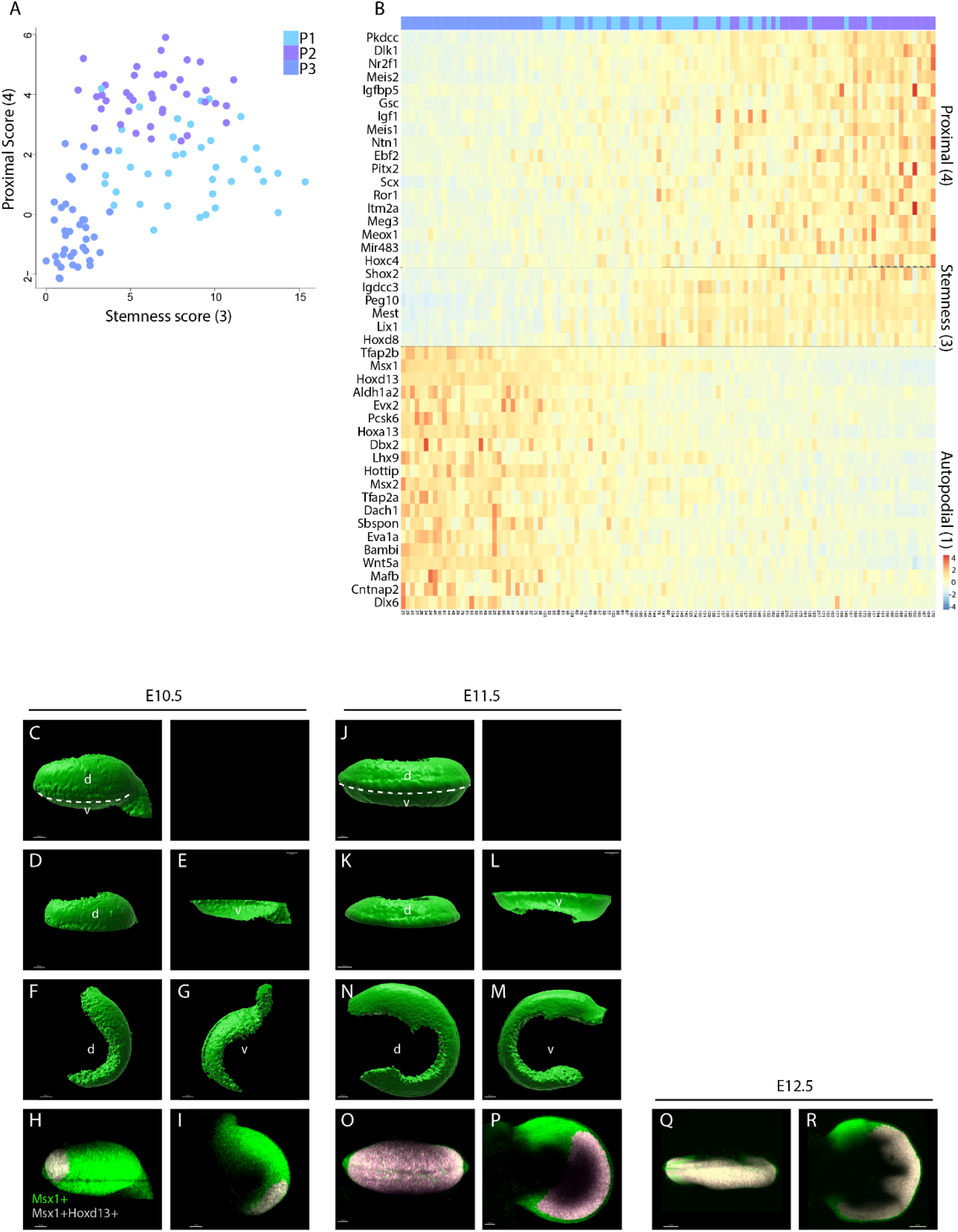
Characterization of mesenchymal progenitor cells. Related to Figure 2. (A) Scatter plot showing the distribution of stemness (*X*-axis) and proximal (*Y*-axis) scores in progenitor metacells. (B) Heatmap showing log_2_ fold change in expression of a subset of proximal, autopodial and stemness module genes in progenitor metacells. (C-G) 3D-segmented and rendered views of the outer surface of *Msx1* mRNA expression domain in E10.5 whole-mount forelimb imaged by light sheet microscopy. (C) Frontal view; dashed white line indicates the plane of optical section in D and E. (D) Dorsal and (E) ventral halves of *Msx1* expression domain show that it extends dorsally and ventrally from the AER midline, and that the dorsal side is wider than the ventral. (F) Dorsal and (G) ventral views show the arc-like shape of Msx1 expression domain and the width differences between the proximal-anterior and proximal-posterior sides. (H,I) Maximum intensity projection (MIP) of *Msx1* and *Hoxd13* mRNA coexpression domain in E10.5 whole-mount forelimb, imaged by light sheet microscopy. (J-N) 3D-segmented and rendered views of the outer surface of *Msx1* mRNA expression domain in E11.5 whole-mount forelimb imaged by light sheet microscopy. (J) Frontal view; dashed white line indicates the plane of optical section in K and L. (K) Dorsal and (L) ventral halves of *Msx1* expression domain show that it extends dorsally and ventrally from the AER midline, and that the dorsal side is wider than the ventral. (N) Dorsal and (M) ventral views show the arc-like shape of *Msx1* expression domain and the width differences between proximal-anterior and proximal-posterior sides. (O-R) MIP of *Msx1* and *Hoxd13* mRNA coexpression domain in E11.5 (O,P) and E12.5 (Q,R) whole-mount forelimbs, imaged by light sheet microscopy. d, dorsal; v, ventral.

**Figure S3.**
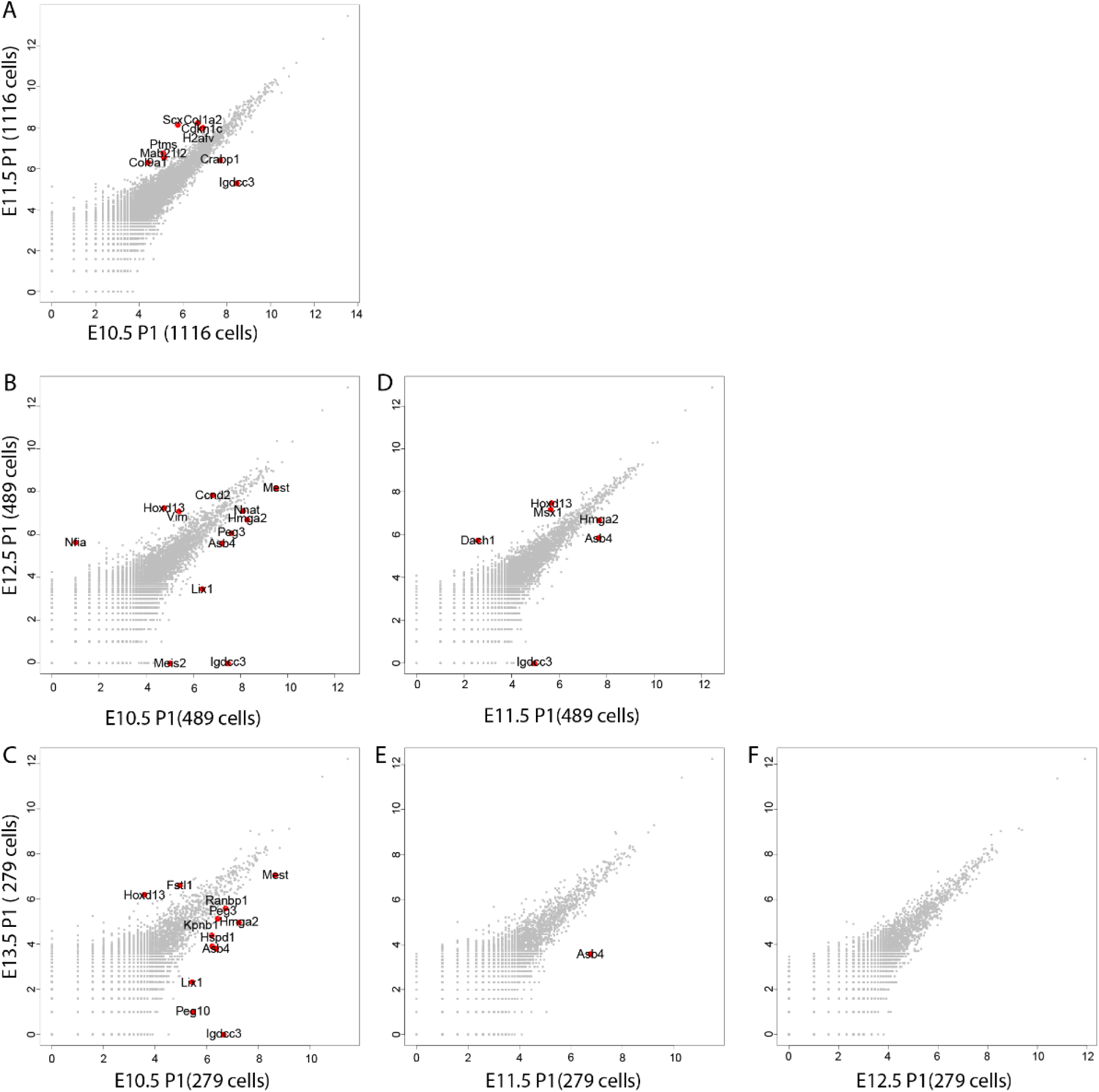
*Msx1* marks the naïve progenitors of the limb. Related to Figure 4. (A-F) Scatter plots showing differentially expressed genes between E10.5-E11.5 (A), E10.5-E12.5 (B), E10.5-E13.5 (C), E11.5-E12.5(D), E11.5-E13.5(E) and E12.5-E13.5 (F) P1 cells. In each comparison, cells were down-sampled for equal cell numbers. Genes that passed a threshold of p-value < 0.01 and log_2_ fold change > 1 in differential expression analysis are indicated by red dots.

**Figure S4.**
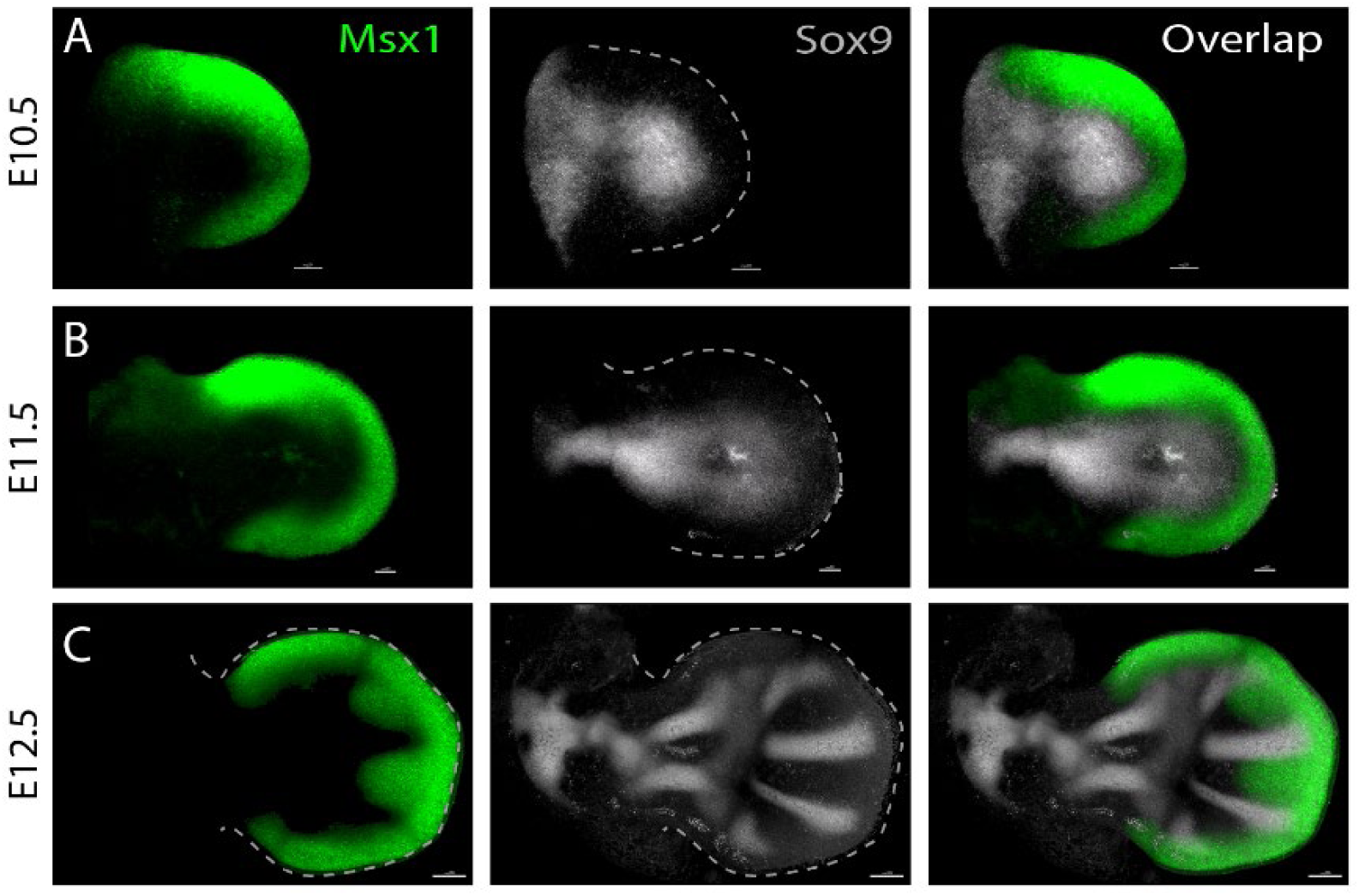
Spatiotemporal analysis of the differentiation of *Msx1^+^* progenitors into *Sox9^+^* cells. Related to Figure 5. (A-C) *In vivo* validation of the opposite trend of *Msx1* and *Sox9* expression, as suggested by the single-cell data. MIP images of E10.5 (A), E11.5 (B) and E12.5 (C) whole-mount forelimbs were stained for *Msx1* and *Sox9* mRNA using *in situ* HCR and imaged by light sheet microscopy. At each stage, n=2.

## Notes

https://www.ncbi.nlm.nih.gov/geo/query/acc.cgi?acc=GSE185940

### Competing Interest Statement

The authors have declared no competing interest.

## References

Akiyama‡, H., Kim‡, J.-E., Nakashima, K., Balmes, G., Iwai, N., Deng, J.M., Zhang, Z., Martin, J.F., Behringer, R.R., Nakamura, T., et al. (2005). Osteo-chondroprogenitor cells are derived from Sox9 expressing precursors. Proc. Natl. Acad. Sci. 102, 14665–14670.

Akiyama, H., Chaboissier, M.-C., Martin, J.F., Schedl, A., and Crombrugghe, B. de (2002a). The transcription factor Sox9 has essential roles in successive steps of the chondrocyte differentiation pathway and is required for expression of Sox5 and Sox6. Genes Dev. 16, 2813–2828.

Akiyama, H., Chaboissier, M.-C., Martin, J.F., Schedl, A., and de Crombrugghe, B. (2002b). The transcription factor Sox9 has essential roles in successive steps of the chondrocyte differentiation pathway and is required for expression of Sox5 and Sox6. Genes Dev. 16, 2813–2828.

Alappat, S., Zhang, Z.Y., and Chen, Y.P. (2003). Msx homeobox gene family and craniofacial development. Cell Res. 2003 136 13, 429–442.

Baran, Y., Bercovich, A., Sebe-Pedros, A., Lubling, Y., Giladi, A., Chomsky, E., Meir, Z., Hoichman, M., Lifshitz, A., and Tanay, A. (2019). MetaCell: analysis of single-cell RNA-seq data using K-nn graph partitions. Genome Biol. 20, 206.

Bell, D.M., Leung, K.K.H., Wheatley, S.C., Ng, L.J., Zhou, S., Ling, K.W., Sham, M.H., Koopman, P., Tam, P.P.L., and Cheah, K.S.E. (1997). SOX9 directly regulates the type-ll collagen gene. Nat. Genet. 1997 162 16, 174–178.

Benoit de Crombrugghe U, Veronique Lefebvre, R.R.B., and Weimin Bi, Shunichi Murakami, W.H. (2000). Transcriptional mechanisms of chondrocyte differentiation. Matrix Biol.

Bensoussan-Trigano, V., Lallemand, Y., Cloment, C. Saint, and Robert, B. (2011). Msx1 and Msx2 in limb mesenchyme modulate digit number and identity. Dev. Dyn. 240, 1190–1202.

Bi, W., Deng, J.M., Zhang, Z., Behringer, R.R., and de Crombrugghe, B. (1999). Sox9 is required for cartilage formation. Nat. Genet. 22, 85–89.

Blitz, E., Sharir, A., Akiyama, H., and Zelzer, E. (2013). Tendon-bone attachment unit is formed modularly by a distinct pool of *Scx* - and *Sox9* -positive progenitors. Development 140, 2680–2690.

Campbell, A.L., Shih, H.-P., Xu, J., Gross, M.K., and Kioussi, C. (2012). Regulation of Motility of Myogenic Cells in Filling Limb Muscle Anlagen by Pitx2. PLoS One 7, e35822.

Capdevila, J., Tsukui, T., Esteban, C.R., Zappavigna, V., and Belmonte, J.C.I. (1999). Control of Vertebrate Limb Outgrowth by the Proximal Factor Meis2 and Distal Antagonism of BMPs by Gremlin. Mol. Cell 4, 839–849.

Chan, H.Y., V., S., Xing, X., Kraus, P., Yap, S.P., Ng, P., Lim, S.L., and Lufkin, T. (2011). Comparison of IRES and F2A-Based Locus-Specific Multicistronic Expression in Stable Mouse Lines. PLoS One 6, e28885.

Chang, H.-M., Martinez, N.J., Thornton, J.E., Hagan, J.P., Nguyen, K.D., and Gregory, R.I. (2012). Trim71 cooperates with microRNAs to repress Cdkn1a expression and promote embryonic stem cell proliferation. Nat. Commun. 2012 31 3, 1–10.

Coudert, A.E., Pibouin, L., Vi-Fane, B., Thomas, B.L., Macdougall, M., Choudhury, A., Robert, B., Sharpe, P.T., Berdal, A., and Lezot, F. (2005). Expression and regulation of the Msx1 natural antisense transcript during development. Nucleic Acids Res. 33, 5208.

Delgado, I., and Torres, M. (2017). Coordination of limb development by crosstalk among axial patterning pathways. Dev. Biol. 429, 382–386.

Delgado, I., López-Delgado, A.C., Roselló-Díez, A., Giovinazzo, G., Cadenas, V., Fernández-de-Manuel, L., Sánchez-Cabo, F., Anderson, M.J., Lewandoski, M., and Torres, M. (2020). Proximo-distal positional information encoded by an Fgf-regulated gradient of homeodomain transcription factors in the vertebrate limb. Sci. Adv. 6, eaaz0742.

Desanlis, I., Paul, R., and Kmita, M. (2020a). Transcriptional Trajectories in Mouse Limb Buds Reveal the Transition from Anterior-Posterior to Proximal-Distal Patterning at Early Limb Bud Stage. J. Dev. Biol. 8, 1–16.

Desanlis, I., Kherdjemil, Y., Mayran, A., Bouklouch, Y., Gentile, C., Sheth, R., Zeller, R., Drouin, J., and Kmita, M. (2020b). HOX13-dependent chromatin accessibility underlies the transition towards the digit development program. Nat. Commun. 2020 111 11, 1–10.

Dudley, A.T., Ros, M.A., and Tabin, C.J. (2002). A re-examination of proximodistal patterning during vertebrate limb development. Nat. 2002 4186897 418, 539–544.

Eyal, S., Kult, S., Rubin, S., Krief, S., Felsenthal, N., Pineault, K.M., Leshkowitz, D., Salame, T.M., Addadi, Y., Wellik, D.M., et al. (2019). Bone morphology is regulated modularly by global and regional genetic programs. Dev. 146.

Fabre, P.J., Leleu, M., Mascrez, B., Lo Giudice, Q., Cobb, J., and Duboule, D. (2018). Heterogeneous combinatorial expression of Hoxd genes in single cells during limb development. BMC Biol. 2018 161 16, 1–15.

Feregrino, C., Sacher, F., Parnas, O., and Tschopp, P. (2019). A single-cell transcriptomic atlas of the developing chicken limb. BMC Genomics 2019 201 20, 1–15.

Fromental-Ramain, C., Warot, X., Messadecq, N., LeMeur, M., Dolle, P., and Chambon, P. (1996). Hoxa-13 and Hoxd-13 play a crucial role in the patterning of the limb autopod. Development 122, 2997–3011.

Huang, A.H. (2017). Coordinated development of the limb musculoskeletal system: Tendon and muscle patterning and integration with the skeleton. Dev. Biol. 429, 420.

Jaitin, D.A., Kenigsberg, E., Keren-Shaul, H., Elefant, N., Paul, F., Zaretsky, I., Mildner, A., Cohen, N., Jung, S., Tanay, A., et al. (2014). Massively parallel single-cell RNA-seq for marker-free decomposition of tissues into cell types. Science (80-.). 343, 776–779.

Johnson, R.L., and Tabin, C.J. (1997). Molecular Models for Vertebrate Limb Development. Cell 90, 979–990.

Kelly, N.H., Huynh, N.P.T., and Guilak, F. (2020). Single cell RNA-sequencing reveals cellular heterogeneity and trajectories of lineage specification during murine embryonic limb development. Matrix Biol. 89, 1.

Kim, D., Langmead, B., and Salzberg, S.L. (2015). HISAT: a fast spliced aligner with low memory requirements. Nat. Methods 12, 357–360.

Knezevic, V., De Santo, R., Schughart, K., Huffstadt, U., Chiang, C., Mahon, K.A., and Mackem, S. (1997). Hoxd-12 differentially affects preaxial and postaxial chondrogenic branches in the limb and regulates Sonic hedgehog in a positive feedback loop. Development 124, 4523–4536.

Lallemand, Y., Nicola, M.-A., Ramos, C., Bach, A., Cloment, C. Saint, and Robert, B. (2005). Analysis of Msx1; Msx2 double mutants reveals multiple roles for Msx genes in limb development. Development 132, 3003–3014.

Lallemand, Y., Bensoussan, V., Cloment, C. Saint, and Robert, B. (2009). Msx genes are important apoptosis effectors downstream of the Shh/Gli3 pathway in the limb. Dev. Biol. 331, 189–198.

Lallemand, Y., Moreau, J., Cloment, C. Saint, Vives, F.L., and Robert, B. (2013). Generation and characterization of a tamoxifen inducible Msx1CreERT2 knock-in allele. Genesis 51, 110–119.

Lefebvre, V.R., Huang, W., Harley, V.R., Goodfellow, P.N., and De Crombrugghe, B. (1997). SOX9 Is a Potent Activator of the Chondrocyte-Specific Enhancer of the Pro␣ 1(II) Collagen Gene. 17, 2336–2346.

Li, D., Sakuma, R., Vakili, N.A., Mo, R., Puviindran, V., Deimling, S., Zhang, X., Hopyan, S., and Hui, C. chung (2014). Formation of Proximal and Anterior Limb Skeleton Requires Early Function of Irx3 and Irx5 and Is Negatively Regulated by Shh Signaling. Dev. Cell 29, 233–240.

Liu, H., Xu, J., Liu, C.-F., Lan, Y., Wylie, C., and Jiang, R. (2015). Whole transcriptome expression profiling of mouse limb tendon development by using RNA-seq. J. Orthop. Res. 33, 840.

Madisen, L., Zwingman, T.A., Sunkin, S.M., Oh, S.W., Zariwala, H.A., Gu, H., Ng, L.L., Palmiter, R.D., Hawrylycz, M.J., Jones, A.R., et al. (2010). A robust and high-throughput Cre reporting and characterization system for the whole mouse brain. Nat. Neurosci. 13, 133–140.

Mariani, F. V., Ahn, C.P., and Martin, G.R. (2008). Genetic evidence that FGFs play an instructive role in limb proximal-distal patterning. Nature 453, 401.

Melton, C., Judson, R.L., and Blelloch, R. (2010). Opposing microRNA families regulate self-renewal in mouse embryonic stem cells. Nat. 2010 4637281 463, 621–626.

Mercader, N., Leonardo, E., Azplazu, N., Serrano, A., Morata, G., Martínez-A, C., and Torres, M. (1999a). Conserved regulation of proximodistal limb axis development by Meis1/Hth. Nat. 1999 4026760 402, 425–429.

Mercader, N., Leonardo, E., Azpiazu, N., Serrano, A., Morata, G., Martínez-A, C., and Torres, M. (1999b). Conserved regulation of proximodistal limb axis development by Meis1/Hth. Nat. 1999 4026760 402, 425–429.

Mercader, N., Leonardo, E., Piedra, M.E., Martinez-A, C., Ros, M.A., and Torres, M. (2000). Opposing RA and FGF signals control proximodistal vertebrate limb development through regulation of Meis genes. Development 127.

Mercader, N., Selleri, L., Criado, L.M., Pallares, P., Parras, C., Cleary, M.L., and Torres, M. (2009). Ectopic Meis1 expression in the mouse limb bud alters P-D patterning in a Pbx1-independent manner. Int. J. Dev. Biol. 53, 1483–1494.

Nagakura, R., Yamamoto, M., Jeong, J., Hinata, N., Katori, Y., Chang, W.-J., and Abe, S. Switching of Sox9 expression during musculoskeletal system development.

Nakamura, Y., Yamamoto, K., He, X., Otsuki, B., Kim, Y., Murao, H., Soeda, T., Tsumaki, N., Deng, J.M., Zhang, Z., et al. (2011). Wwp2 is essential for palatogenesis mediated by the interaction between Sox9 and mediator subunit 25. Nat. Commun. 2, 251.

Nassari, S., Duprez, D., and Fournier-Thibault, C. (2017a). Non-myogenic Contribution to Muscle Development and Homeostasis: The Role of Connective Tissues. Front. Cell Dev. Biol. 0, 22.

Nassari, S., Blavet, C., Bonnin, M.-A., Stricker, S., Duprez, D., and Fournier-Thibault, C. (2017b). The chemokines CXCL12 and CXCL14 differentially regulate connective tissue markers during limb development. Sci. Rep. 7, 17279.

Niswander, L. (2003). Pattern formation: old models out on a limb. Nat. Rev. Genet. 2003 42 4, 133–143.

Pastor, W.A., Liu, W., Chen, D., Ho, J., Kim, R., Hunt, T.J., Lukianchikov, A., Liu, X., Polo, J.M., Jacobsen, S.E., et al. (2018). TFAP2C regulates transcription in human naive pluripotency by opening enhancers. Nat. Cell Biol. 2018 205 20, 553–564.

Patterson, M., Chan, D.N., Ha, I., Case, D., Cui, Y., Handel, B. Van, Mikkola, H.K., and Lowry, W.E. (2011). Defining the nature of human pluripotent stem cell progeny. Cell Res. 2011 221 22, 178–193.

Pellegrini, M., Pantano, S., Fumi, M.P., Lucchini, F., and Forabosco, A. (2001). Agenesis of the Scapula in Emx2 Homozygous Mutants. Dev. Biol. 232, 149–156.

Petit, F., Sears, K.E., and Ahituv, N. (2017). Limb development: a paradigm of gene regulation. Nat. Rev. Genet. 2017 184 18, 245–258.

Probst, S., Kraemer, C., Demougin, P., Sheth, R., Martin, G.R., Shiratori, H., Hamada, H., Iber, D., Zeller, R., and Zuniga, A. (2011). SHH propagates distal limb bud development by enhancing CYP26B1-mediated retinoic acid clearance via AER-FGF signalling. Development 138, 1913–1923.

Pryce, B.A., Brent, A.E., Murchison, N.D., Tabin, C.J., and Schweitzer, R. (2007). Generation of transgenic tendon reporters, ScxGFP and ScxAP, using regulatory elements of the scleraxis gene. Dev. Dyn. 236, 1677–1682.

Reijntjes, S., Stricker, S., and Mankoo, B.S. (2007). A comparative analysis of Meox1 and Meox2 in the developing somites and limbs of the chick embryo. Int. J. Dev. Biol.

Rybak, A., Fuchs, H., Hadian, K., Smirnova, L., Wulczyn, E.A., Michel, G., Nitsch, R., Krappmann, D., and Wulczyn, F.G. (2009). The let-7 target gene mouse lin-41 is a stem cell specific E3 ubiquitin ligase for the miRNA pathway protein Ago2. Nat. Cell Biol. 2009 1112 11, 1411–1420.

Salbaum, J.M. (1998). Punc, a novel mouse gene of the immunoglobulin superfamily, is expressed predominantly in the developing nervous system. Mech. Dev. 71, 201–204.

Saunders, J.W. (1948). The proximo-distal sequence of origin of the parts of the chick wing and the role of the ectoderm. J. Exp. Zool. 108, 363–403.

Schweitzer, R., Chyung, J.H., Murtaugh, L.C., Brent, A.E., Rosen, V., Olson, E.N., Lassar, A., and Tabin, C.J. (2001). Analysis of the tendon cell fate using Scleraxis, a specific marker for tendons and ligaments. Development 128.

Scotti, M., Kherdjemil, Y., Roux, M., and Kmita, M. (2015a). A Hoxa13: Cre Mouse Strain for Conditional Gene Manipulation in Developing Limb, Hindgut, and Urogenital System. Genesis 53, 366.

Scotti, M., Kherdjemil, Y., Roux, M., and Kmita, M. (2015b). A Hoxa13: Cre Mouse Strain for Conditional Gene Manipulation in Developing Limb, Hindgut, and Urogenital System. Genesis 53, 366.

Shen, H., Wilke, T., Ashique, A.M., Narvey, M., Zerucha, T., Savino, E., Williams, T., and Richman, J.M. (1997). Chicken Transcription Factor AP-2: Cloning, Expression and Its Role in Outgrowth of Facial Prominences and Limb Buds. Dev. Biol. 188, 248–266.

Sheth, R., Bastida, M.F., and Ros, M. (2007). Hoxd and Gli3 interactions modulate digit number in the amniote limb. Dev. Biol. 310, 430–441.

Soeda, T., Deng, J.M., de Crombrugghe, B., Behringer, R.R., Nakamura, T., and Akiyama, H. (2010). Sox9-expressing precursors are the cellular origin of the cruciate ligament of the knee joint and the limb tendons. Genesis 48, 635–644.

Stricker, S., Mathia, S., Haupt, J., Seemann, P., Meier, J., and Mundlos, S. (2012). Odd-Skipped Related Genes Regulate Differentiation of Embryonic Limb Mesenchyme and Bone Marrow Mesenchymal Stromal Cells. Stem Cells Dev. 21, 623–633.

Sugimoto, Y., Takimoto, A., Akiyama, H., Kist, R., Scherer, G., Nakamura, T., Hiraki, Y., and Shukunami, C. (2013). Scx+/Sox9+ progenitors contribute to the establishment of the junction between cartilage and tendon/ligament. Development 140, 2280–2288.

Summerbell, D., Lewis, J.H., and Wolpert, L. (1973). Positional Information in Chick Limb Morphogenesis. Nature 244, 492–496.

Sun, C., Zhang, T., Liu, C., Gu, S., and Chen, Y. (2013). Generation of Shox2-Cre allele for tissue specific manipulation of genes in the developing heart, palate, and limb. Genesis 51, 515.

Sun, X., Mariani, F. V., and Martin, G.R. (2002). Functions of FGF signalling from the apical ectodermal ridge in limb development. Nat. 2002 4186897 418, 501–508.

Tabin, C., and Wolpert, L. (2007a). Rethinking the proximodistal axis of the vertebrate limb in the molecular era. Genes Dev. 21, 1433–1442.

Tabin, C., and Wolpert, L. (2007b). Rethinking the proximodistal axis of the vertebrate limb in the molecular era. Genes Dev. 21, 1433–1442.

Tickle, C. (2005). Making digit patterns in the vertebrate limb. Nat. Rev. Mol. Cell Biol. 2006 71 7, 45–53.

Treweek, J.B., Chan, K.Y., Flytzanis, N.C., Yang, B., Deverman, B.E., Greenbaum, A., Lignell, A., Xiao, C., Cai, L., Ladinsky, M.S., et al. (2015). Whole-body tissue stabilization and selective extractions via tissue-hydrogel hybrids for high-resolution intact circuit mapping and phenotyping. Nat. Protoc. 2015 1011 10, 1860–1896.

Tzchori, I., Day, T.F., Carolan, P.J., Zhao, Y., Wassif, C.A., Li, L., Lewandoski, M., Gorivodsky, M., Love, P.E., Porter, F.D., et al. (2009). LIM homeobox transcription factors integrate signaling events that control three-dimensional limb patterning and growth. Development 136, 1375–1385.

Ulrike Hirning-Folz, Monika Wilda, Volkhard Rippe, Jorn Bullerdiek, and H.H. (1998). The expression pattern of the Hmgic gene during development. Genes Chromosom. Cancer 23, 350–357.

Vallecillo-García, P., Orgeur, M., Vom Hofe-Schneider, S., Stumm, J., Kappert, V., Ibrahim, D.M., Börno, S.T., Hayashi, S., Relaix, F., Hildebrandt, K., et al. (2017). Odd skipped-related 1 identifies a population of embryonic fibro-adipogenic progenitors regulating myogenesis during limb development. Nat. Commun.

Vickerman, L., Neufeld, S., and Cobb, J. (2011). Shox2 function couples neural, muscular and skeletal development in the proximal forelimb. Dev. Biol. 350, 323–336.

Wang, J., Rao, S., Chu, J., Shen, X., Levasseur, D.N., Theunissen, T.W., and Orkin, S.H. (2006). A protein interaction network for pluripotency of embryonic stem cells. Nat. 2006 4447117 444, 364–368.

Watson, B.A., Feenstra, J.M., Arsdale, J.M. Van, Rai-Bhatti, K.S., Kim, D.J.H., Coggins, A.S., Mattison, G.L., Yoo, S., Steinman, E.D., Pira, C.U., et al. (2018). LHX2 Mediates the FGF-to-SHH Regulatory Loop during Limb Development. J. Dev. Biol. 2018, Vol. 6, Page 13 6, 13.

Wilson-Rawls, J., Rhee, J.M., and Rawls, A. (2004). Paraxis Is a Basic Helix-Loop-Helix Protein That Positively Regulates Transcription through Binding to Specific E-box Elements *. J. Biol. Chem. 279, 37685–37692.

Wolpert, L. (2002). The progress zone model for specifying positional information. Int. J. Dev. Biol. 46, 869–870.

Worringer, K.A., Rand, T.A., Hayashi, Y., Sami, S., Takahashi, K., Tanabe, K., Narita, M., Srivastava, D., and Yamanaka, S. (2014). The let-7/LIN-41 Pathway Regulates Reprogramming to Human Induced Pluripotent Stem Cells by Controlling Expression of Prodifferentiation Genes. Cell Stem Cell 14, 40–52.

Xianjin Zhou, Kathleen F. Benson, H.R.A.& K.C. (1995). Mutation responsible for the mouse pygmy phenotype in the developmentally regulated factor HMGI-C. Nature 376, 771–774.

Yang, Y., and Wilson, M.J. (2015). Lhx9 gene expression during early limb development in mice requires the FGF signalling pathway. Gene Expr. Patterns 19, 45–51.

Yang, B., Treweek, J.B., Kulkarni, R.P., Deverman, B.E., Chen, C.-K., Lubeck, E., Shah, S., Cai, L., and Gradinaru, V. (2014). Single-Cell Phenotyping within Transparent Intact Tissue through Whole-Body Clearing. Cell 158, 945–958.

Yu, J., Vodyanik, M.A., Smuga-Otto, K., Antosiewicz-Bourget, J., Frane, J.L., Tian, S., Nie, J., Jonsdottir, G.A., Ruotti, V., Stewart, R., et al. (2007). Induced Pluripotent Stem Cell Lines Derived from Human Somatic Cells. Science (80-.). 318, 1917–1920.

Zákány, J., Fromental-Ramain, C., Warot, X., and Duboule, D. (1997). Regulation of number and size of digits by posterior Hox genes: A dose-dependent mechanism with potential evolutionary implications. Proc. Natl. Acad. Sci. 94, 13695–13700.

Zeller, R., López-Ríos, J., and Zuniga, A. (2009). Vertebrate limb bud development: moving towards integrative analysis of organogenesis. Nat. Rev. Genet. 10, 845–858.

Zhang, J., Tam, W.-L., Tong, G.Q., Wu, Q., Chan, H.-Y., Soh, B.-S., Lou, Y., Yang, J., Ma, Y., Chai, L., et al. (2006). Sall4 modulates embryonic stem cell pluripotency and early embryonic development by the transcriptional regulation of Pou5f1. Nat. Cell Biol. 2006 810 8, 1114–1123.

Zhang, J., Ratanasirintrawoot, S., Chandrasekaran, S., Li, H., Collins, J.J., and Daley Correspondence, G.Q. (2016). LIN28 Regulates Stem Cell Metabolism and Conversion to Primed Pluripotency Accession Numbers GSE67568. Cell Stem Cell 19, 66–80.

Zhao, F., Bosserhoff, A.-K., Buettner, R., and Moser, M. (2011). A Heart-Hand Syndrome Gene: Tfap2b Plays a Critical Role in the Development and Remodeling of Mouse Ductus Arteriosus and Limb Patterning. PLoS One 6, e22908.

Zuniga, A. (2015). Next generation limb development and evolution: old questions, new perspectives. Development 142, 3810–3820.

